# HSV Tegument Protein pUL51 Interacts with Host Dynactin for Viral Spread

**DOI:** 10.1101/2022.03.21.485087

**Authors:** Shaowen White, My Tran, Richard J. Roller

**Affiliations:** Department of Microbiology, University of Iowa, Iowa City, IA 52242; Department of Biological Sciences, Southern Illinois University Edwardsville, Edwardsville, IL 62026

**Keywords:** HSV-1, UL51, peptide inhibitor, anti-viral drug, dynactin, viral spread

## Abstract

Dynein motors are microtubule associated protein complexes that mediate multiple essential cellular processes, such as long-distance cargo trafficking and stabilization of the microtubule organization center. Most of these functions and their regulations depend on the dynein motor subunit dynactin. By using an infection-inducible system, we disrupted dynein motor function after HSV entry by overexpressing a dominant-negative inhibitor of dynein, resulting in a 5-fold growth defect in Vero cells and 1000-fold growth defect in CAD neuronal cells. Also, we found that in infected CAD cells, the dynein complex was recruited to viral assembly sites regardless of microtubule polymerization. Based on these observations, we then identified a novel interaction between conserved HSV-1 tegument protein pUL51 and p150Glued. pUL51 is a palmitoylated Golgi membrane-associated protein that is required for efficient virus assembly and spread. Overexpression of pUL51 alone was sufficient to recruit p150Glued to Golgi membranes. Sequences that are important and sufficient for pUL51-p150Glued interaction were mapped to residues 90 to 124 in pUL51 and residues 548 to 911 in p150Glued. Deletion of a.a 90-124 in pUL51 resulted in a moderate viral growth defect, a profound spread defect, and failure to accumulate both dynactin and the viral spread factor glycoprotein E (gE) at cell-cell junctions. A synthetic peptide that contains pUL51 a.a 90-125 could also inhibit viral growth and spread in pUL51-dependent manner. Taken together, our results suggest that the proper function of pUL51 in efficient viral assembly and spread depends on its interaction with p150Glued.

**Author Summary:** During their life-long infections, herpesviruses can spread within the human body in spite of a powerful immune response that includes antibodies that inactivate the virus. The virus accomplishes this by targeting its newly made infectious particles to junctions where infected cells meet their uninfected neighbors. Junctional targeting of virus particles requires that the virus hijack normal cellular machinery used for moving large cargoes within the cell, a process that is only partially understood. In this study, we have discovered a critical interaction between a virus protein that is present in all herpesviruses and a host cell cargo-moving protein that is required for virus spread between cells and shown that this interaction is common to several medically important human herpesviruses. We have also identified a short piece of this virus protein that is responsible for the interaction and shown that this short piece can be used to interrupt the interaction and prevent spread of herpes simplex virus (HSV) in cultured cells. Our study provides prove-of-concept evidence that the spread pathway of HSV, and possibly other herpesviruses, can be targeted for therapeutic purposes.

## Introduction

Herpesviruses are large DNA viruses that are of significant global health concern, especially in immune-compromised populations including neonates and tissue transplant patients (1, 2). Currently, the only licensed therapeutics for treating herpesvirus infection are acyclovir and its derivatives, which all target a single viral mechanism - the viral DNA polymerase. This poses a concern over the rise of drug resistance (3).

There are ample potential alternative drug targets for a pan-herpesvirus inhibitor, as there are at least 18 conserved genes among all three subfamilies of herpesviruses that are not directly related to DNA replication (4). These genes are essential for the viral life cycle in vivo, and many of them play roles in viral assembly and egress. Some of these conserved assembly proteins participate in self-assembly complexes, such as major and minor capsid proteins; while others, including the viral nuclear egress complex, the virus protein kinases pUS3 and pUL13, pUL21, pUL36, VP26, pUS9, and gE interact with host proteins to mediate other aspects of viral egress (5–13). Targeting these viral-host interactions for therapeutics may reduce development of drug resistance since the host partner is not subject to selective pressure (14). Currently, many modulators of host factors are being explored for their potential as targets of anti-viral therapy, including ESCRT pathway modulators for HIV (15, 16) and sirtuin activators for HCMV (17), however, little has been explored for these approaches in targeting assembly proteins of herpes simplex virus (HSV).

The assembly and egress of herpesviruses can be divided into five steps: (i) the assembly and packaging of DNA into viral capsids inside the nucleus, (ii) the transportation of viral capsids across nuclear membranes into the cytoplasm, (iii) the transportation of cytoplasmic capsids and cellular membranes that carry virus envelope and tegument proteins towards the cytoplasmic assembly site(s), (iv) capsid budding into the cytoplasmic envelopment membranes to form mature virions, and (v) sorting and transportation of mature virions outward, either towards apical surface for apical release or towards cell-cell junctions for cell-cell spread (CCS) (18). Steps iii and v extensively require the involvement of microtubule-dependent trafficking mechanisms, as naked capsids, vesicles containing viral proteins, and vesicles containing mature virions are too large to move by diffusion (18). The dynein involvement in trafficking of egressing capsids is evidenced by the observation that capsids accumulate in the nucleus upon the deletion of inner tegument protein pUL36, which is required for dynein-capsid interaction (10, 19, 20). Dynein involvement in glycoprotein transport is implied by the presence of functional cytoplasmic retrieval motifs on the cytoplasmic tail of some glycoproteins such as gE and gB (13, 21). These retrieval motifs interact with clathrin adaptor proteins to mediate endocytosis, of which downstream transportation requires microtubule-associated motors, such as dynein (22, 23).

How microtubule-dependent motors function in trafficking of mature virions is incompletely understood. This step is essential for HSV CCS *in vivo*, which allows progeny virions to infect adjacent cells without exposing to neutralizing antibodies in the environment (24), or to transmit between epithelial cells and neurons (25, 26). Genetic evidence in HSV and HCMV suggests that herpesvirus CCS is a specialized pathway distinct from apical release (27–29). In HSV, pUL51, gE, and pUS9 are proteins that are known to mediate CCS, with pUS9 being proposed to be a kinesin (microtubule plus end-directed motor) adaptor in neuronal cells (12, 30–32). Since trafficking for release and spread occurs immediately after the assembly of progeny virions, factors that play role in this step are likely also concentrated at the assembly site(s).

Among HSV proteins that are involved in viral assembly and CCS, tegument protein pUL51 is thought to play critical roles, since deletion of pUL51 results in as large as 100-fold defect in viral production and ∼40-fold defect in CCS (33, 34). pUL51 is also considered as one of the herpesvirus “core genes”, as its homologs are present in all families of herpesviruses (4). pUL51 and its homologs have similar functions (35–38) and conserved viral interaction partners (39), suggesting that pUL51 and its homologs function in similar ways. Thus, pUL51 function is an attractive pan-herpesvirus therapeutic target.

Dynein motor function is required for formation and function of virus assembly sites in neuronal cells. We also find that HSV-1 pUL51 interacts with the host dynein adaptor dynactin major subunit p150Glued and that this interaction facilitates virus assembly in neuronal cells and is required for efficient spread in epithelial cells. We also generated pUL51-derived recombinant peptides that inhibit HSV CCS and growth in pUL51-dependent manner while showing no cytotoxicity in cell culture. Lastly, we show preliminary data that the interaction between pUL51 and p150Glued might be conserved among all subfamilies of herpesvirus.

## Results

### Disrupting dynein function results in a HSV assembly defect

We have shown that HSV forms a cytoplasmic assembly center (cVAC) in several neuronal cells, including CathA (CAD) cells (35). Since HSV cVAC formation is microtubule- and dynein-dependent, we hypothesize that the dynein motor complex is involved in viral assembly. To test this hypothesis, we specifically disrupted dynein function by overexpressing the CC1 domain of the dynactin subunit p150Glued in HSV-infected cells. Overexpression of the CC1 domain has been shown to disrupt dynein motor function by interfering with dynactin/dynein complex formation (40, 41). Since it has previously been shown that initiation of infection by HSV is dynein-dependent (42),the effect of dynein motor function disruption was limited to late steps in infection by using plasmids in which the EGFP-CC1 domain fusion protein (abbreviated as CC1) or EGFP expression was under the control of the promotor of HSV gene UL34, which leads to infection-inducible expression of these proteins (Fig 1A) (43). Overexpression of CC1 prevented formation or maintenance of HSV cVAC as shown by dispersed staining for the virion envelope protein, gE (Fig 1 B and C). Some leaky expression was observed for both EGFP and CC1 in uninfected cells (Fig 1A), but this was not sufficient to inhibit initiation of infection, since the frequency of infected cells was not diminished in cells expressing CC1 compared to those expressing EGFP (Supplemental Fig 1A). Overexpression of CC1 also led to a ∼2-fold viral single-step growth defect in transfected cell cultures (Fig 1D). Since only ∼50% of infected cells were expressing CC1 (Fig 1E), this suggested that dynein disruption might severely inhibit viral assembly. To test this possibility, we constructed infection-inducible plasmids that not only express EGFP or CC1, but also express the major capsid protein VP5. CAD and Vero cells were transfected with these plasmids and then infected with VP5-null HSV, so that viral progeny could only be produced in cells that over-express EGFP- or CC1 and VP5. The EGFP control and CC1-expressing plasmids were transfected at the same efficiency (Supplemental Fig 1, C and D), but CC1 overexpression diminished viral replication more than 1000-fold in CAD cells (Fig 1F) and ∼5-fold in Vero cells (Fig 1G), suggesting that dynein plays important roles in efficient HSV assembly. The virus assembly function of dynein in CAD cells was not due to an effect on capsid nuclear egress, since accumulation of fluorescently tagged capsids in the cytoplasm was unaffected by CC1 overexpression in CAD cells (Supplemental Fig 2).

**Fig 1.**
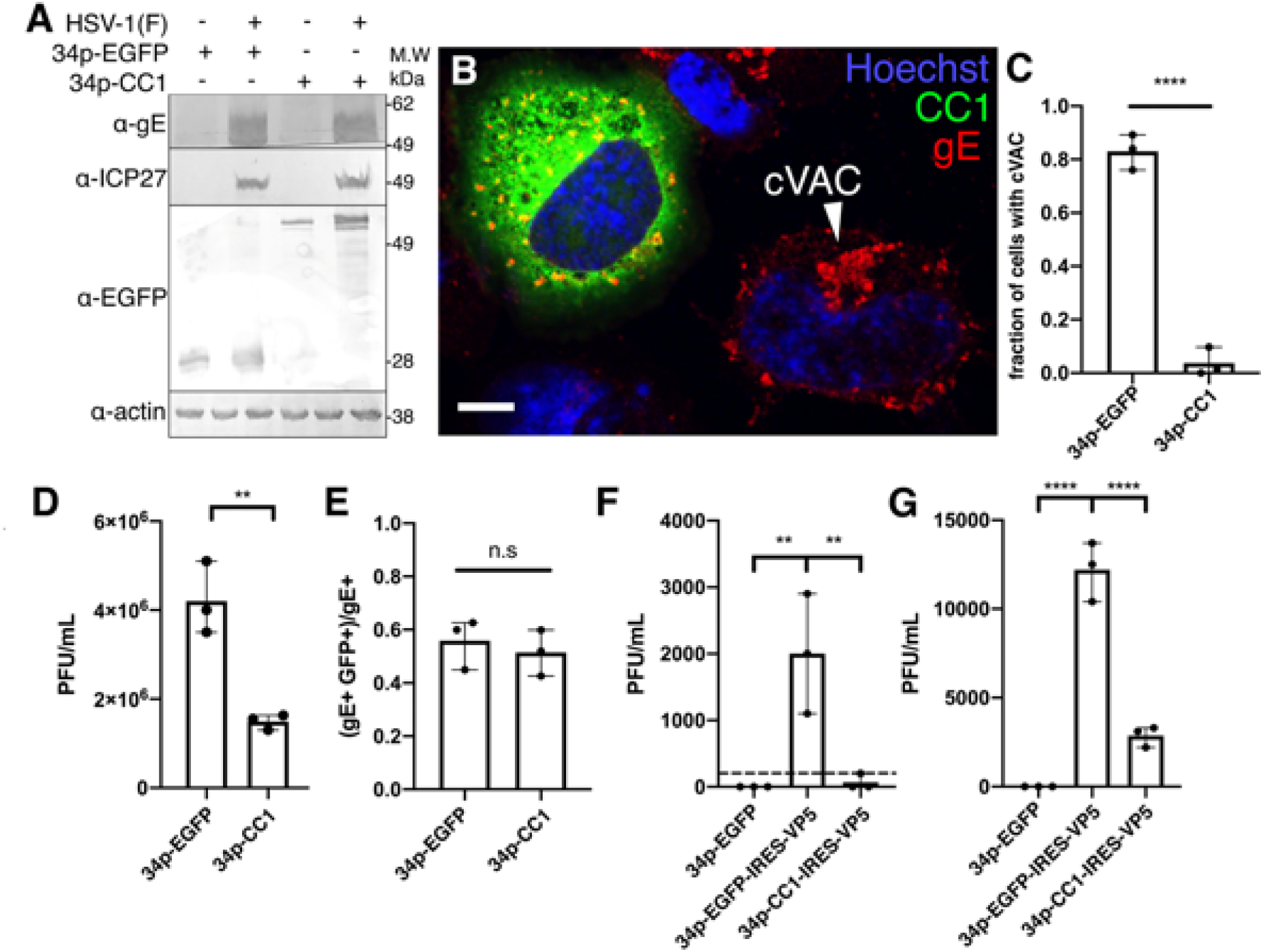
Disrupting dynein function by CC1 overexpression impedes viral assembly. (A) Immunoblots showing HSV-1 infection-induced, UL34 promoter-controlled expression of EGFP from plasmid 34p-GFP, or CC1 from plasmid 34p-CC1 in CAD cells. (B) Confocal image of 34p-CC1-transfected, HSV-1-infected CAD cells. Nuclei are stained with Hoechst33342 in blue; CC1-EGFP is shown in green and the location of glycoprotein gE is shown in red. The white arrowhead indicates the cVAC in the un-transfected cell. Scale bar represents 10µm. (C) Fraction of EGFP or CC1 and gE double-positive CAD cells that form a cVAC in confocal images obtained 12 hours post-infection (h.p.i.). At least 50 cells were imaged and scored for presence of a cVAC in three independent experiments. (D) Viral production in transfected-superinfected CAD cells at 16 hr after infection at m.o.i.=5. (E) Fraction of infected CAD cells that express EGFP or CC1 in confocal images obtained 12 h.p.i. At least 104 infected (i.e., gE-expressing cells) were imaged at 12 h.p.i. and scored for expression of EGFP or CC1 in three independent experiments. (F, G) Viral production in CAD (F) or Vero (G) cells after transfection for 24 hr with indicated plasmid and then infection for an additional 18 hr with VP5-null virus at m.o.i.=5. Progeny virion production was quantification by plaque assay on VP5-complementing cells. In panels (C-G), **=P<0.01, ***=P<0.001, ****=P<0.0001, data represent mean ± RNG.

### Dynein and dynactin concentrate to the HSV cVAC

Since the dynein motor complex apparently functions in formation or maintenance of the HSV-1 assembly center, the localization of dynein and dynactin components of the complex was investigated in CAD cells. In uninfected CAD cells, both dynein intermediate chain (DIC) and p150Glued were concentrated at the microtubule organization center (MTOC) (Fig 2A and B). In infected cells, most DIC was concentrated at the cVAC, where it co-localized with the HSV structural protein pUL11 (Fig 2C, arrows). In contrast, p150Glued localized both at the cVAC and as intense, small puncta throughout the cytoplasm outside the cVAC (referred to here as “peripheral puncta”). Many of these peripheral p150Glued puncta did not contain detectable DIC (Fig 2C) and did not represent fragmented microtubules (Fig 2D). These puncta did, however, partially coincide with peripheral gE and pUL11 puncta (Fig 2E, arrows), and with some viral capsids (Fig 2F, yellow arrows).

**Fig 2.**
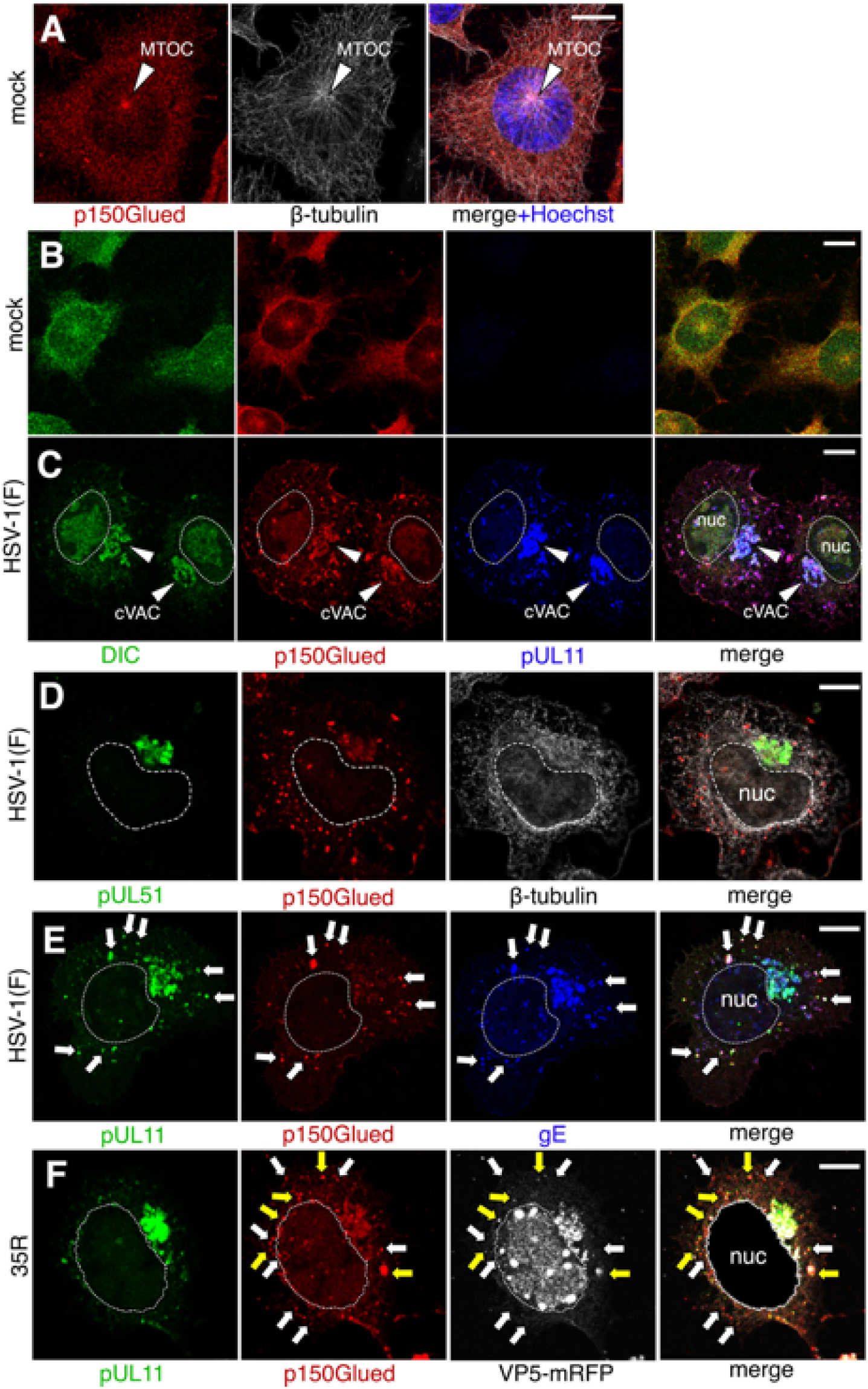
Localization of DIC and p150Glued in infected CAD cells. (A, B) Mock infected CAD cells that were immunofluorescently stained for indicated proteins. (C-F) CAD cells infected with HSV (C-E) or HSV-1 35R at m.o.i=5 for 12 hr before fixation and staining. All scale bars represent 10µm. In (E), arrows indicate puncta outside the cVAC where pUL11, p150Glued, and gE colocalize. In (F), white arrows indicate p150Glued puncta outside the cVAC that did not contain detectable capsid signal, while yellow arrows indicate p150Glued puncta outside the cVAC that colocalize with capsids.

The co-localization of DIC and p150Glued at the cVAC with virus membrane-bound structural proteins pUL51 and pUL11 suggested association between the dynein motor complex and cytoplasmic membranes. Since p150Glued can interact with vesicular cargos, we hypothesized that HSV induced membrane association of p150Glued. As shown in Fig 3, in vehicle treated CAD cells, p150Glued and the Golgi apparatus both concentrated around the MTOC. Upon treatment with nocodazole, a drug that prevents microtubule polymerization, p150Glued became cytoplasmic, and did not co-localize with dispersed Golgi membranes (Fig 3B). In contrast, in HSV infected cells, nocodazole treatment dispersed p150Glued-positive structure, but these structures remained punctate and were still associated both with virus structural proteins and with Golgi membranes (Fig 3D, arrows). These results suggest that HSV infection causes nocodazole-resistant association between p150Glued and viral assembly compartment membranes.

**Fig 3.**
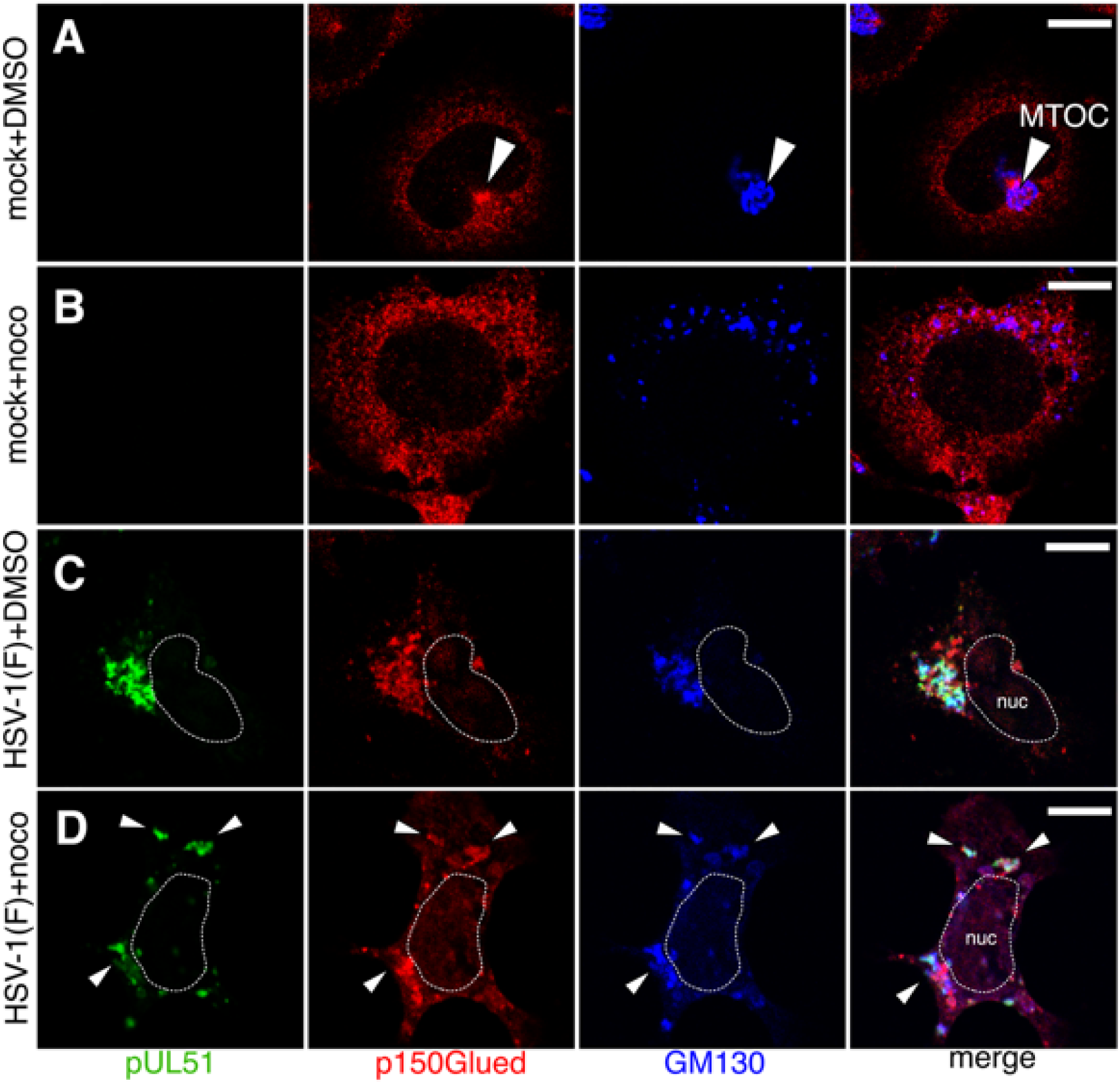
p150Glued association with Golgi membranes is nocodazole-resistant in infected cells. (A, B) CAD cells were treated with DMSO (A) or 5µg/ml nocodazole (B) for 9 hr. (C, D) CAD cells were infected with HSV at m.o.i=5 for 12 hr before fixation and staining. At 3 h.p.i, culture media were replaced with media containing DMSO (C) or 5µg/ml nocodazole (D). Arrows in (D) indicate clusters of Golgi membranes that are associated with p150Glued puncta. All scale bars represent 10µm.

### Dynactin interacts with HSV pUL51

The membrane association of p150Glued is mediated by protein-protein interactions (23) and we hypothesized that a viral structural protein might contribute to the membrane association of p150Glued in infected cells. It has previously been shown that overexpressed EGFP-p150Glued localizes to microtubules and induces the formation of microtubule bundles (44). Expression of this fusion protein also disrupts dynein motor function and results in dispersal of the Golgi apparatus into puncta that are distributed throughout the cytoplasm. We used the microtubule bundling property of overexpressed EGFP-p150Glued to look for viral proteins that are recruited to these microtubule bundles in infected Vero cells. We observed that in EGFP-p150Glued-overexpressing HSV-infected cells, a fraction of pUL51 colocalized with filamentous EGFP-p150Glued instead of Golgi marker GM130, while this filamentous localization was not observed for pUL11, gD, and gE (Fig 4, B and C). The recruitment of pUL51 away from Golgi membranes and onto microtubule bundles by p150Glued could also be achieved in co-transfected cells, without the presence of other viral proteins (Fig 5B). Also, overexpressing pUL51 alone was sufficient to recruit the native form of p150Glued to membranes (Fig 5C), suggesting pUL51 might associate with p150Glued directly or in complex with other host factors. The p150Glued-pUL51 interaction was not specific to Vero cells and could also be observed HEp2 cells (human epithelial-derived), CAD cells, and SH-SY5Y cells (human neuroblastoma-derived) (Fig 5 D-F).

**Fig 4.**
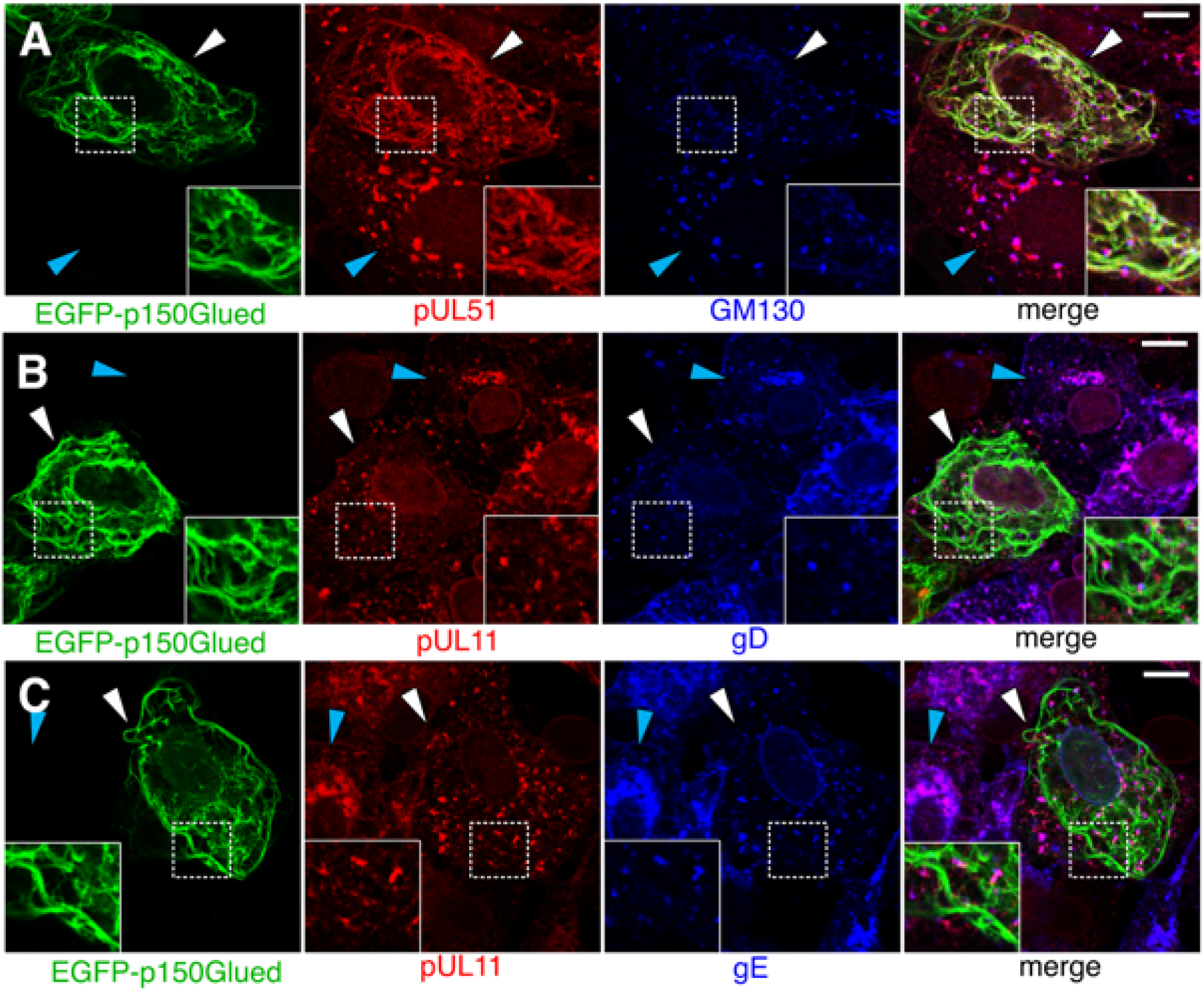
Overexpressed EGFP-p150Glued recruits pUL51 to microtubule bundles in infected Vero cells. Cells were transfected with EGFP-p150Glued plasmid for 24 hr before infection with HSV for 14 hr. White arrowheads indicate cells that express EGFP-p150Glued. Blue arrowheads indicate untransfected cells. Insets show zoomed images in dotted-line boxes. Scale bars represent 10µm.

**Fig 5.**
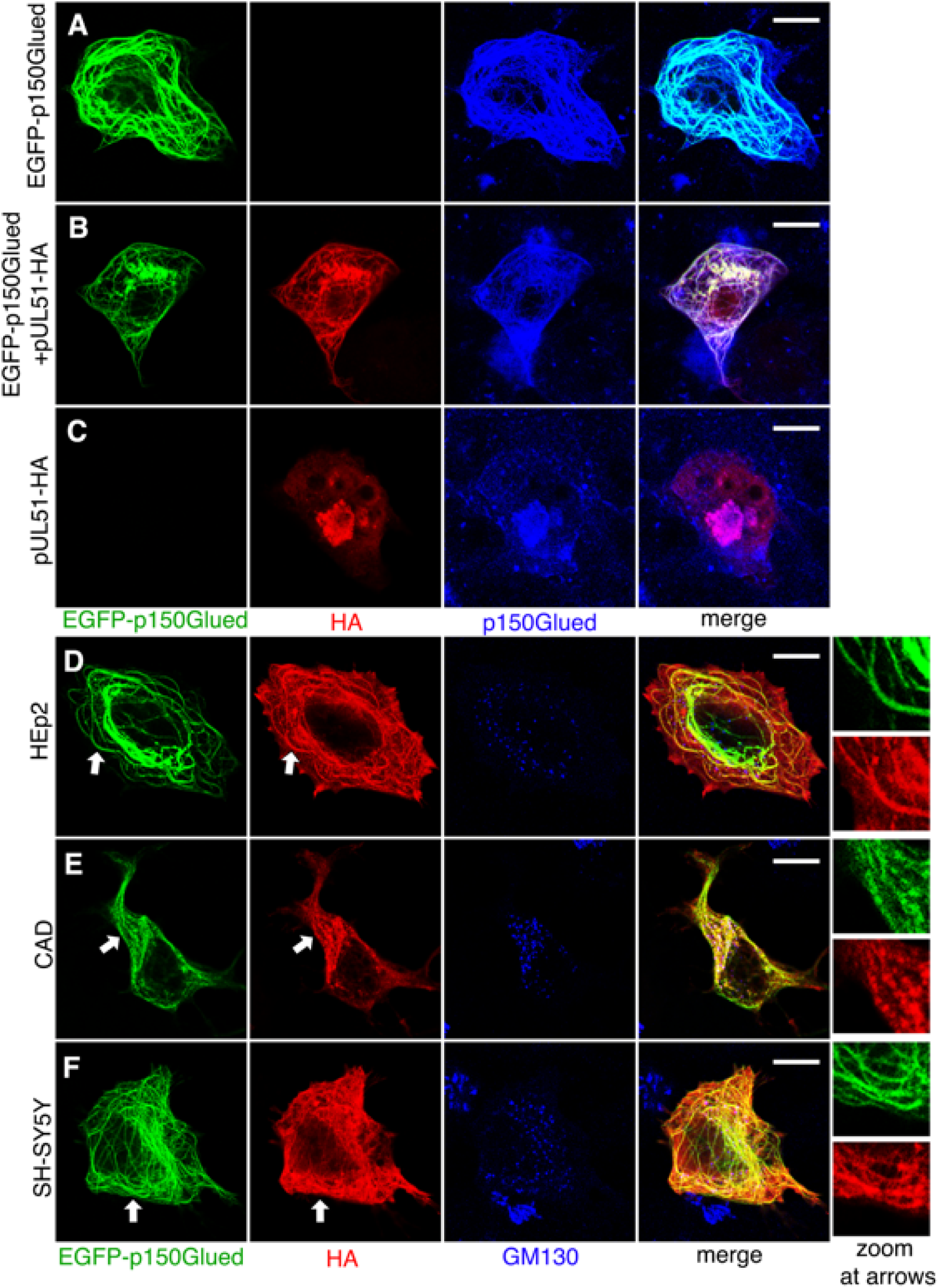
Overexpressed EGFP-p150Glued recruits pUL51 to microtubule bundles in co-transfected cells. (A-C) Single (A, C) or double (B) transfected Vero cells were stained for pUL51-HA and p150Glued. Note that the brightness of p150Glued staining in (C) has been increased to match signal intensity of overexpressed EGFP-p150Glued in (A) and (B). (D-F) Indicated cells were co-transfected with EGFP-p150Glued and pUL51-HA and immunofluorescently stained using anti-HA and anti-GM130. Scale bars represent 10µm.

Since EGFP-p150Glued might cause a microtubule structure abnormality (44), and since pUL51 associates with membranes by palmitoylation of a N-terminal cystine residue (45), we tested whether p150Glued-pUL51 interaction requires microtubule polymerization or pUL51 membrane-association. As shown in Fig 6, neither nocodazole treatment nor mutation of the pUL51 palmitoylated residue prevented EGFP-p150Glued colocalization with pUL51 in co-transfected cells, suggesting that neither is needed for the interaction.

**Fig 6.**
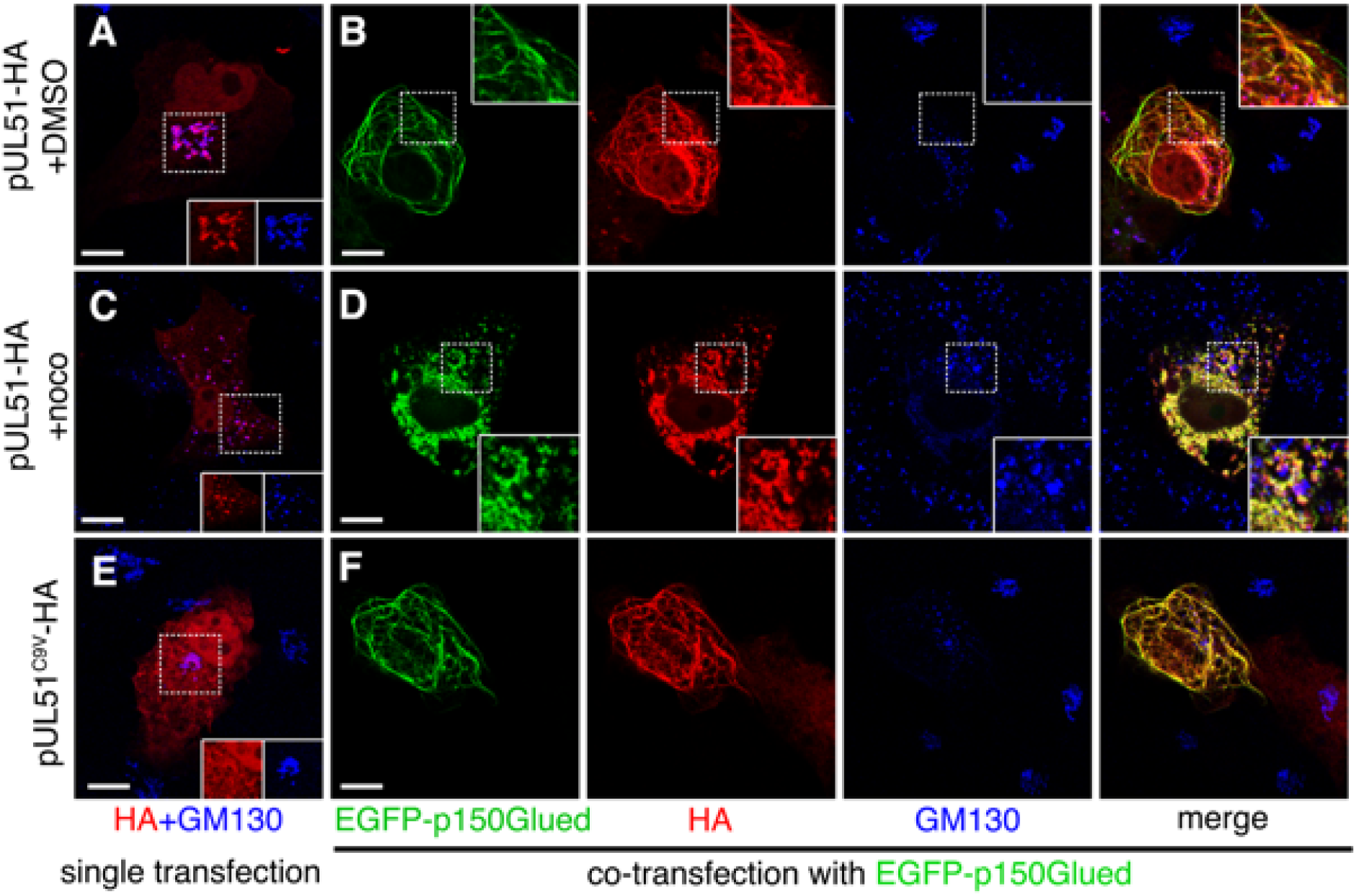
Colocalization of pUL51 and EGFP-p150Glued is independent of microtubule polymerization and pUL51 membrane association. (A-D) Vero cells were transfected with pUL51-HA (A, C) or co-transfected with pUL51-HA and EGFP-p150Glued (B, D). Transfected cells were treated with DMSO (A, B) or nocodazole (C, D) for 2 hr. (E, F) Vero cells were transfected with pUL51^C9V^-HA (E) or co-transfected with pUL51^C9V^-HA and EGFP-p150Glued (F). Insets in (A, C, E) show split channels of images in dotted-line boxes. Insets in (B,D,E) show zoomed images in dotted-line boxes. Scale bars represent 10µm.

### Identifying sequences in pUL51 and p150Glued that are important for their interaction

We next identified sequences in pUL51 that are important for p150Glued-pUL51 interaction by co-transfecting Vero cells with EGFP-p150Glued and various truncations of pUL51 (Fig 7A). pUL51^125-244^ was unable to localize with p150Glued, while pUL51^90-244^ remained filamentous, suggesting a.a 90-125 of pUL51 is necessary for EGFP-p150Glued-pUL51 interaction (Fig 7B). However, pUL51^1-124^ only colocalized with EGFP-p150Glued weakly (Fig 7B, arrows) while pUL51^1-166^ had drastically improved colocalization, suggesting sequences between a.a 124-166 of pUL51 improved the interaction efficiency between pUL51 and p150Glued. The crystal structure of pUL51 in complex with pUL7 has been partially solved and indicates that a.a 90-125 mostly forms an extended alpha helix (Fig 7, C-E). Amino acids 124-166 could not be resolved in the structure, but are predicted to include another extended alpha helix (Fig 7C). We tested if a.a 90-125 is sufficient for p150Glued-pUL51 interaction by tagging mCherry with pUL51^90-125^ and co-expressing it with EGFP-p150Glued in Vero cells. Indeed, pUL51^90-125^-mCherry was sufficient to interact with EGFP-p150Glued in assays for both co-localization (Fig 7F-G) and co-immunoprecipiation (Fig 7J).

**Fig 7.**
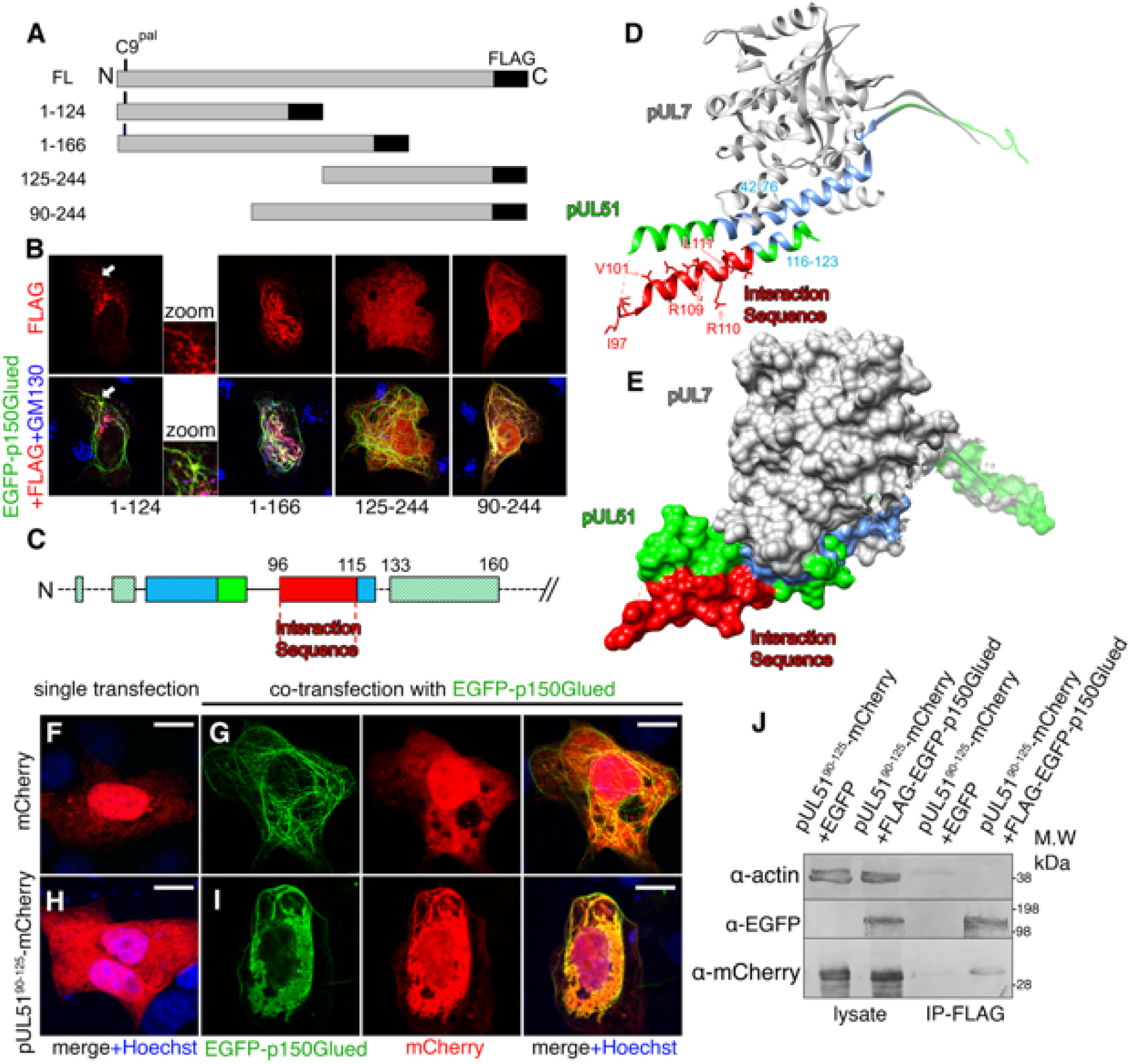
pUL51 a.a 90-125 are sufficient for recruitment by EGFP-p150Glued. (A) Schematic showing various FLAG-tagged pUL51 truncations. The top bar depicts full-length (FL) pUL51. Numbers to the left of each of the lower bars indicates the pUL51 sequences that are present in the truncation construct. Small vertical lines near the N-terminus of first three truncations represent the palmitoylated cystine. (B) Confocal images showing recruitment of pUL51 truncations by EGFP-p150Glued to microtubule bundles in co-transfected Vero cells. Zoomed images display filamentous localization pattern of pUL51^1-124^-FLAG at the locations indicated by arrows. (C) Schematic for secondary structures of pUL51 a.a 1-160. Boxes (green and blue) represent α-helixes. Blue, α-helixes that interact with pUL7. Solid lines and colors represent sequences that have been solved in the crystal structure. Dashed lines and striped color fillings represent secondary structures predicted by NetSurfP-2.0 (76). (D, E) Crystal structure of pUL51 (PDB:6T5A) displayed by secondary structure (D) or surface (E). Color scheme is the same as in (C). (F-I) Confocal images showing Vero cells that were transfected with indicated plasmids. Split EGFP and mCherry channels are shown for (G) and (I). Scale bars represent 10µm. (J) 293T cells were transfected with indicated plasmid combinations. Cell lysates were subjected to IP with anti-FLAG beads.

### The interaction between p150Glued and pUL51 might play roles in viral production

We next tested the functional significance of p150Glued-pUL51 interaction. To this end, we attempted to construct a pUL51 mutant that has impaired p150Glued interaction. Residues 90-125 of pUL51 contain several hydrophobic and charged residues that might be expected to participate in protein-protein interactions (Fig 7D). However, single mutations of several hydrophobic residues (I97A, V101A, L111A), or combined mutation of two neighboring arginine residues (R109A and R110A), or combined mutation of three residues in the hypothetically exposed region on the N-terminus end of region 90-125 (G91A, L92A, E93A) did not disrupt EGFP-p150Glued-pUL51 interaction in co-transfected cells (Supplemental Fig 3). We then deleted a.a 90-125 in pUL51, which indeed resulted in reduced co-immunoprecipitation (Fig 8A) and disrupted co-localization with p150Glued in co-transfected Vero cells (Fig 8, B and C). Surprisingly, whereas WT pUL51-HA is tightly localized to Golgi membranes in transfected cells, in a significant portion of pUL51^d90-125^-HA single-overexpressing Vero cells, pUL51^d90-125^-HA did not localize to Golgi membranes (Fig 8, D and E). A similar phenotype was also observed in transfected CAD cells, although the difference was subtle, and only occurred in less than a half of cells (Fig 8, F and G).

**Fig 8.**
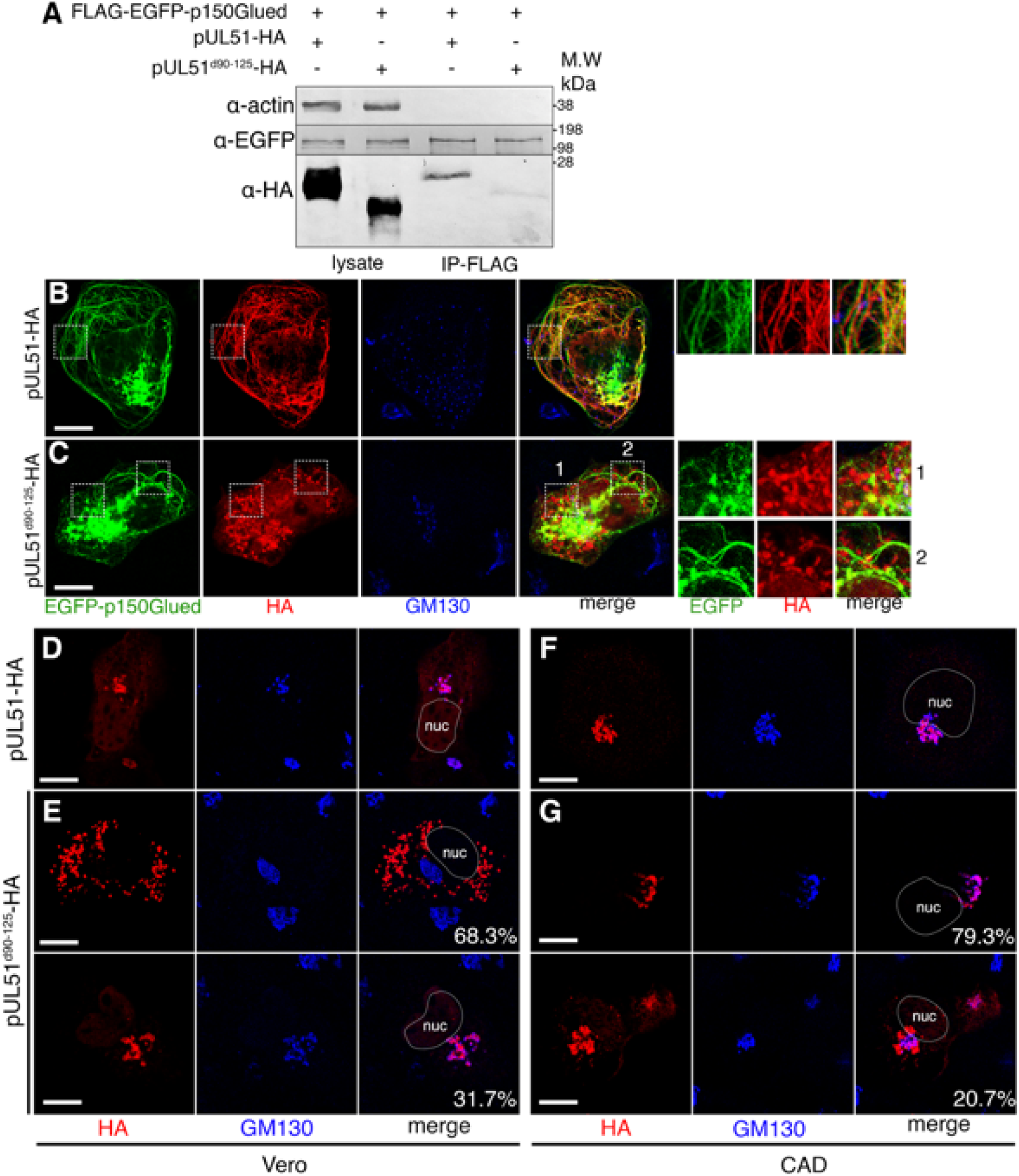
pUL51^d90-125^ mutant protein interacts less efficiently with EGFP-p150Glued and Golgi membranes in transfected cells. (A) 293T cells were transfected with indicated plasmid combinations. Cell lysates were subjected to IP with anti-FLAG beads. (B, C) Confocal images of Vero cells co-transfected with EGFP-p150Glued and WT (B) or mutant (C) HA-tagged pUL51. Right, zoomed images showing EGFP, HA, and merged channels in dotted line boxes. Inset boxes 1 and 2 feature areas of punctate and filamentous p150Glued respectively. (D-G) Vero (D, E) or CAD (F, G) cells were transfected with indicted pUL51 constructs. (E, G) Upper panels display the majority of phenotype and lower panels display the minority. 100 cells were imaged for each condition, and the percentage of cells displaying each phenotype were indicated in the merged images. All scale bars represent 10µm.

To investigate the biological consequences of the loss of pUL51 a.a 90-125, we engineered the UL51^d90-125^ mutation into BAC-derived HSV-1(F) virus. Two isolates were constructed independently to reduce possibility of phenotype arising from unintended mutations. pUL51^d90-125^ had a reduced expression level in comparison to the WT control (Fig 9A), but still localized to Golgi membranes (Fig 9C), suggesting other viral factors also play roles in the Golgi-localization of pUL51. In UL51^d90-125^ infected CAD cells, although the cVAC still formed, its organization was altered so that pUL51 vesicles no longer associated with gE vesicle within the cVAC (Fig 9, D-F). At 16 h.p.i, the d90-125 mutation resulted in ∼10-fold growth defect compared to WT control. The mutant virus also grew ∼10-fold better than a pUL51^d73-244^ mutant, in which most of the UL51 coding sequence had been deleted (Fig 9, G and H) (33). From this we conclude that a.a 90-125 of pUL51 plays role in viral assembly.

**Fig 9.**
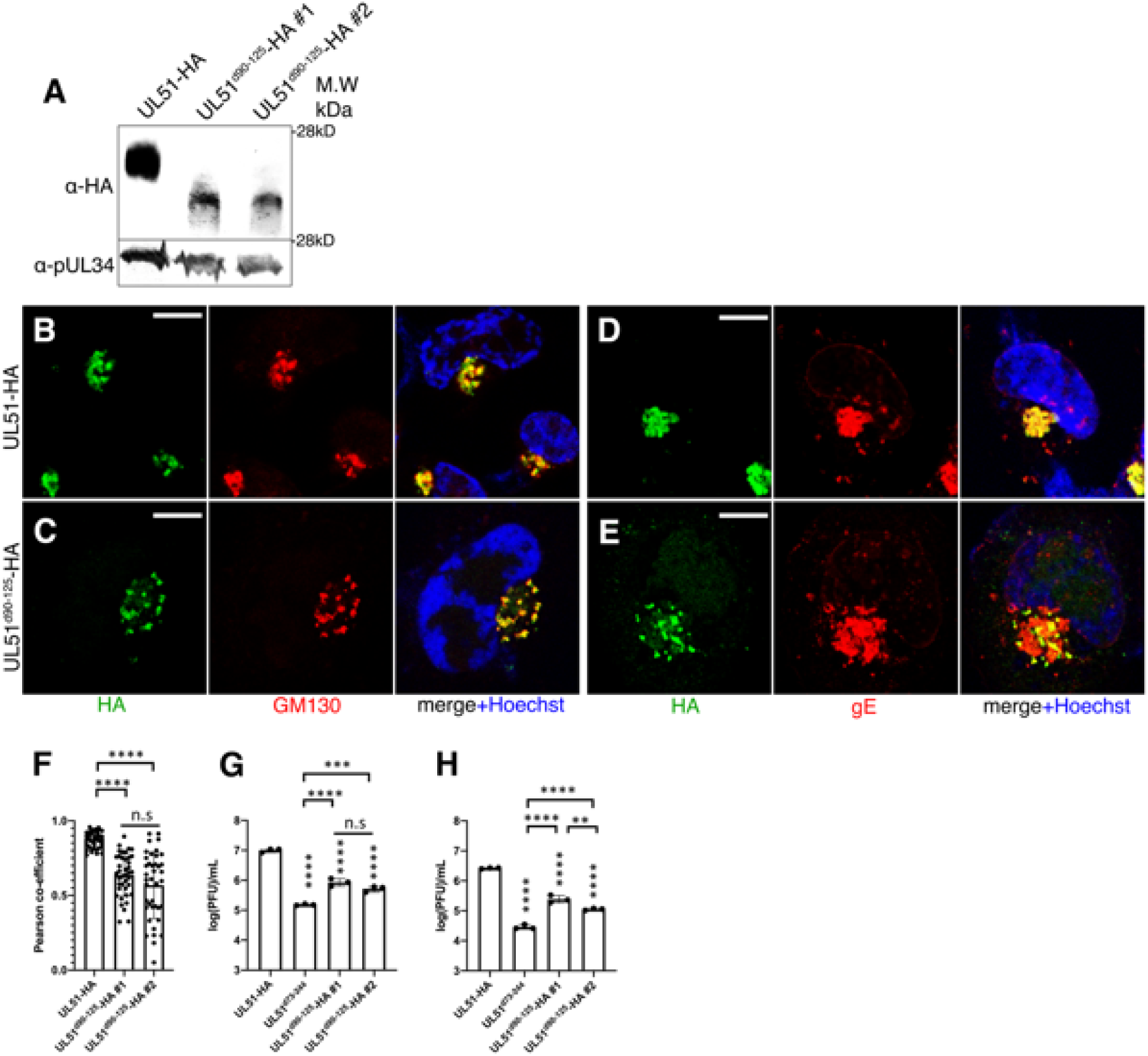
UL51^d90-125^ mutation disrupts cVAC formation and viral production in infected cells. (A) Immunoblot of lysates of CAD cells infected for 16 hr at m.o.i.=5 by viruses that express HA-tagged WT or mutant pUL51. (B-E) Representative confocal images of infected CAD cells at 12 h.p.i. Scale bars represent 10µm. The infecting virus is indicted to the left of the panels. (F) Pearson co-efficient for co-localization between gE and pUL51 in cells represented in (D) and (E). Each point represents one cell. (G, H) Vero (G) or CAD (H) cells were infected at m.o.i=5 by indicated HSV mutants and infectious progeny production was determined at 16 h.p.i. **=P<0.01, ***=P<0.001, ****=P<0.0001. Vertical columns of asterisks indicate the P values for the comparison with UL51-HA. Data represent mean ± SD.

### The interaction between p150Glued and pUL51 might play roles in viral spread and release

Since pUL51 plays important role in cell-cell spread of HSV (33, 34), the viral spread and release phenotype was investigated. As shown in Fig 10, A and B, UL51^d90-125^ mutants had >40-fold defect in the ability to spread on a monolayer of cells in the presence of neutralizing antibody. They also accumulated much less gE at cell-cell junctions late in the infection (Fig 10, C-F). Surprisingly, in infected Vero cells, p150Glued also accumulated at cell-cell junctions (Fig 10C), and this localization also depended on pUL51 a.a 90-125 (Fig 10, D, E, and G). To elucidate whether the spread defect of UL51d90-125 was due to a specific disruption in the CCS pathway or disruption in both apical release and CCS, we measured apical release of UL51 mutant viral particles late in infection. At 30 h.p.i, UL51^d90-125^ mutants had reduced viral production and apical release phenotypes that were indistinguishable from the UL51^d73-244^ mutant (Fig 10, H and I). An apical release defect of the UL51^d90-125^ mutant was also observed in HaCaT cells at 24 h.p.i (Fig 10, J and K).

**Fig 10.**
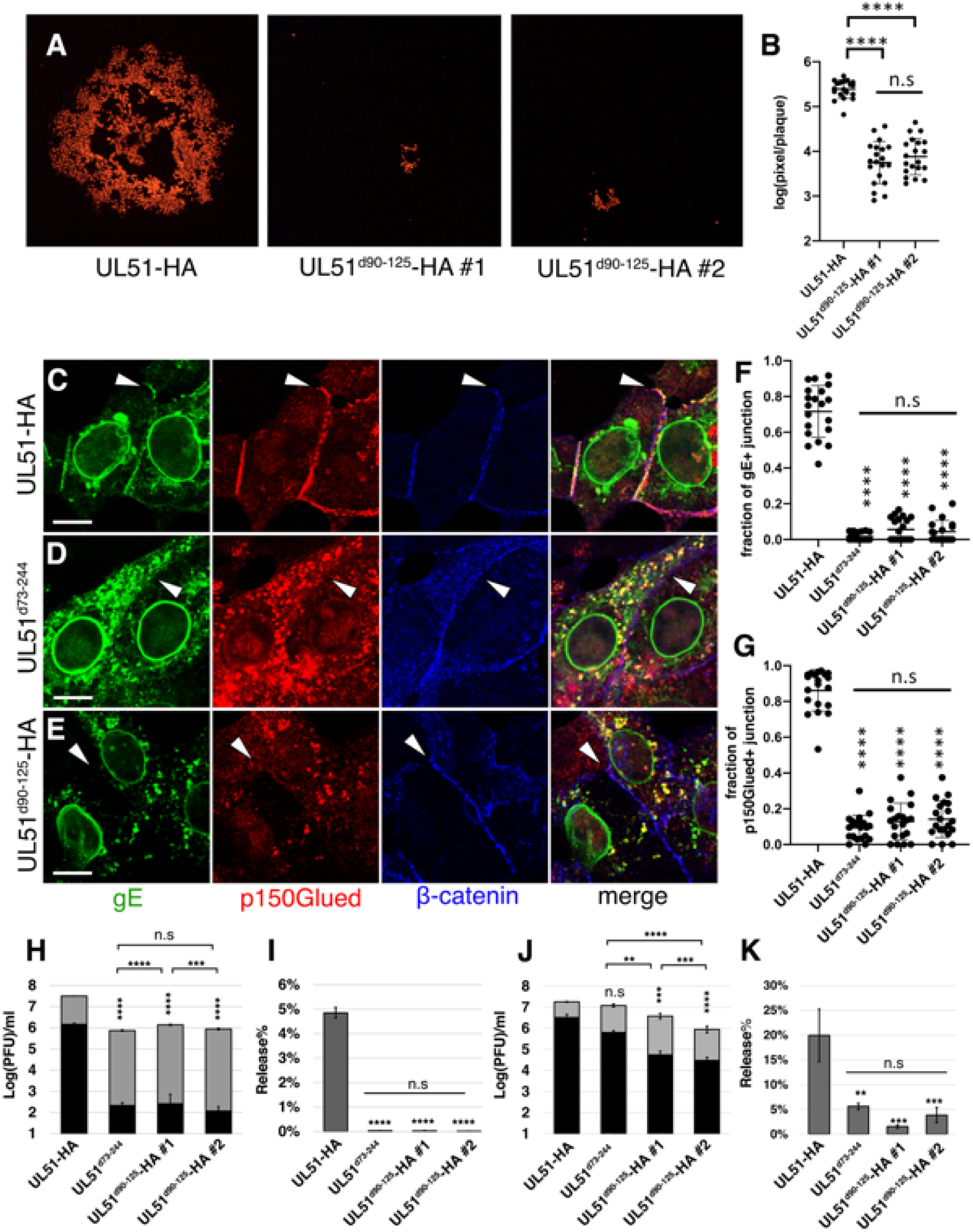
UL51^d90-125^ mutation impedes CCS and viral release. (A, B) Representative images (A) and quantification (B) showing plaque size difference between control and mutant HSV. 100 PFU of each indicated virus were used to infect 9 cm^2^ of a confluent monolayer of Vero cells. Infected cells were incubated in the presence of neutralizing antibody for 72 hr before fixation. pUL26.5 was detected by immunofluorescence and used to indicate infected cells. (C-G) Confocal imaging-based quantification of gE and p150Glued accumulation at cell-cell junctions. Vero cells were infected at m.o.i=5 and fixed 14 h.p.i. Scale bars represent 10µm. Each dot in graphs in (F) and (G) represents the fraction of cell-cell junctions that were also positive for indicated protein in an individual field of view. (H-K) Quantification of released or cell-associated infectious progenies in Vero (H, I) or HaCaT (J, K) cells. Cells were infected at m.o.i=5. Supernatants and cell lysates of infected Vero or HaCaT cells were separated and harvested 30 h.p.i or 24 h.p.i respectively. (H, J) Grey bars in each graph represent the amount of cell-associated infectious progeny. Black bars represent infectious progeny in the supernatant. (I, K) Comparison of viral release efficiency represented by [supernatant PFU]/[cell-associated PFU]. **=P<0.01, ***=P<0.001, ****=P<0.0001. Data represent mean ± SD.

### pUL51 a.a 90-125 is a dominant-negative inhibitor of pUL51 functions

It is possible that the growth and spread defect of UL51^d90-125^ mutants was caused by an abnormality in protein folding. Since UL51^90-125^-mCherry was sufficient to interact with p150Glued (Fig 7), we hypothesized that overexpression of UL51^90-125^-mCherry fusion protein would dominant-negatively inhibit WT pUL51 function. To test this hypothesis, pUL51^90-125^-mCherry was co-expressed with WT pUL51 in CAD cells. Indeed, overexpression of pUL51^90-125^-mCherry was sufficient to reproduce the Golgi-dissociation phenotype of UL51^d90-125^ mutants (Fig 11, A-C). When overexpressed in infected cells, pUL51^90-125^-mCherry resulted in a moderate growth defect, and this growth defect depended on the presence of wild-type pUL51 (Fig 11, D-G), suggesting expression of pUL51 a.a 90-125-mCherry fusion protein interfered with normal pUL51 function.

**Fig 11.**
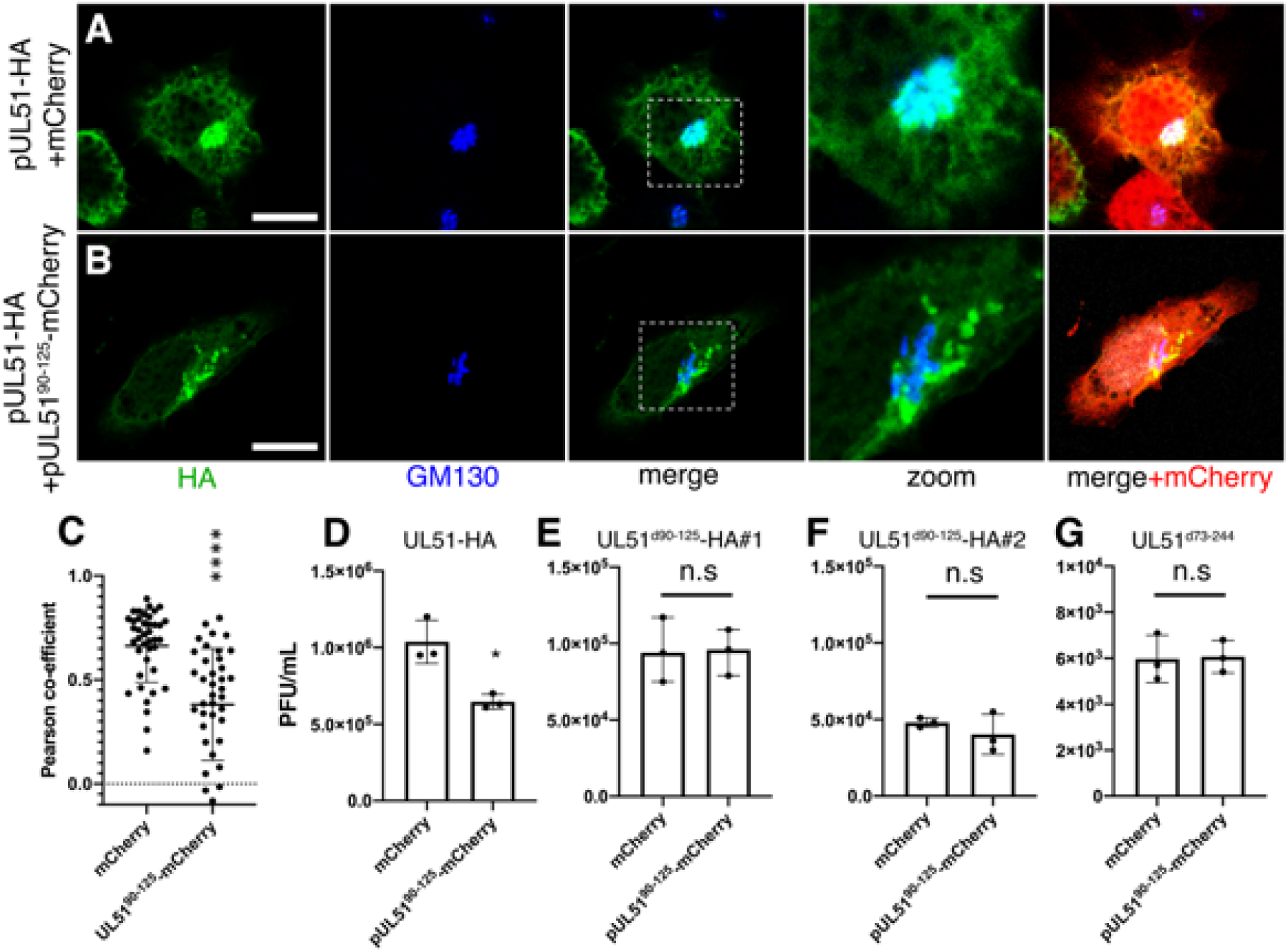
Overexpression of pUL51^90-125^-mCherry fusion protein disrupts normal pUL51 function. (A, B) Representative images of CAD cells that were co-transfected with plasmids expressing the proteins indicated to the left of the panels. Scale bars represent 10µm. (C) Pearson co-efficient analysis for colocalization of GM130 and pUL51 in cells represented in (A) and (B). (D-G) Virus production in CAD cells that were transfected with mCherry or pUL51^90-125^-mCherry 24 hr before infection with the indicated virus at m.o.i=5. Infected cell lysates were harvested at 16 h.p.i. and the amount of infectious progeny was determined by plaque assay on pUL51-complementing cells. *=P<0.05, ****=P<0.0001. Data represent mean ± SD.

To test whether a.a 90-125 of pUL51 alone could interfere with the CCS function of pUL51, we designed peptides containing pUL51 a.a 90-125, with either no tag, or a HIV tat-tag to facilitate membrane penetration (Table 1) (46). At 7nM, none of the peptides significantly impeded cell growth (Fig 12A), and two of the peptides reduced viral production in Vero cells, with C-terminus-tagged peptide 90125tat having the greatest effect (Fig 12C). This inhibition of growth by peptide 90125tat was not observed in UL51^d73^ mutant infected cells (Fig 12D), suggesting the inhibition is pUL51-dependent and, therefore, that the peptide interfered with pUL51 function. At 7nM, tat90125 and 90125tat peptides reduced WT HSV-1(F) BAC virus spread (Fig 12, E and F). To test if this inhibition effect is specific to the HSV-1(F) BAC clone, the same set of experiments was conducted for WT HSV-1(F). As shown in Fig 12, E and G, both tat-tagged peptides reduced WT HSV-1(F) virus spread, with the 90125tat peptide having the most significant effect on plaque size. The effect of 90125tat peptide depended on the presence of wild-type pUL51 (Fig 12 H-J). The 90125tat peptide also reduced the accumulation of gE and p150Glued at cell-cell junctions (Fig 12, K-M).

**Fig 12.**
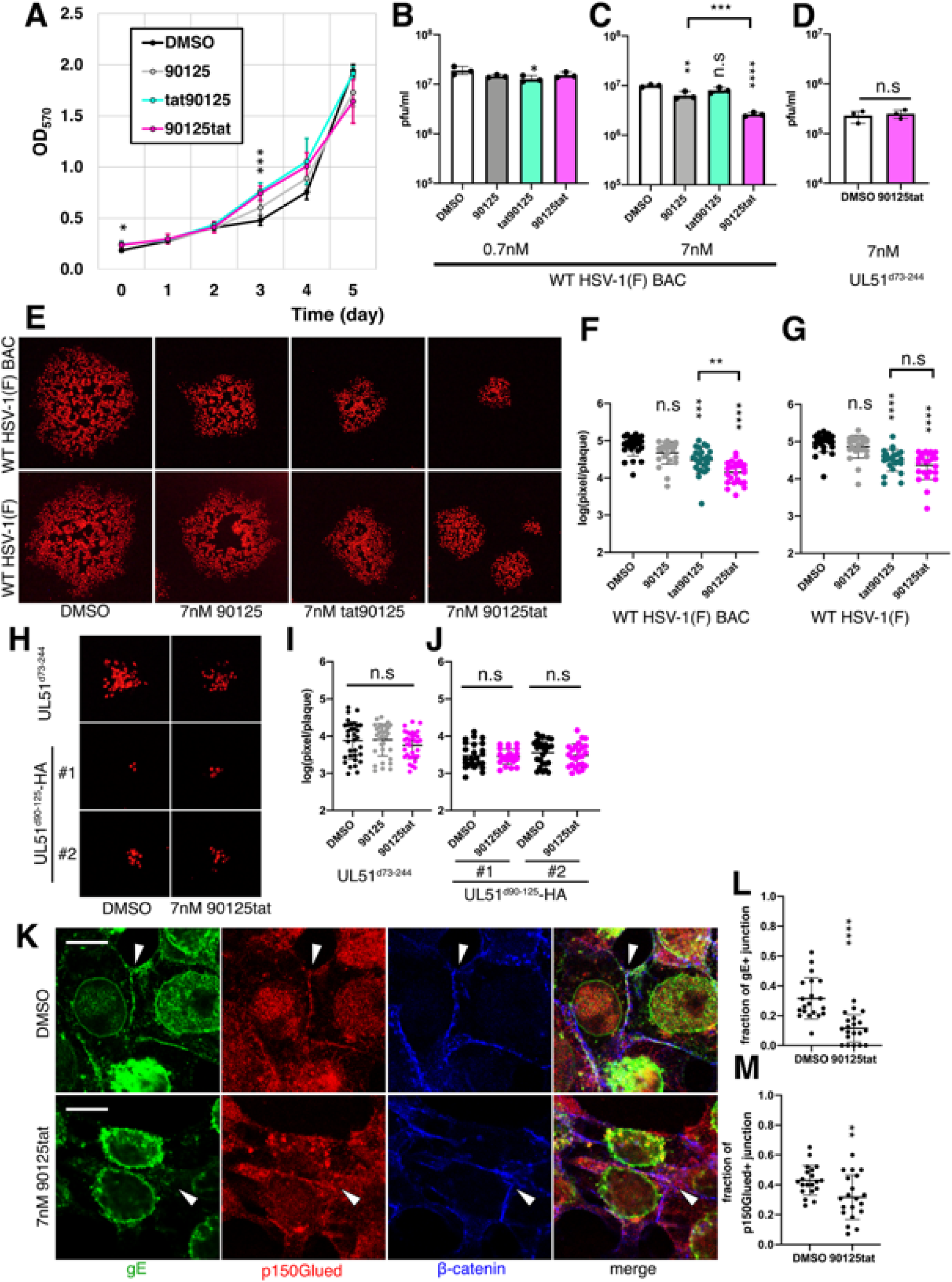
pUL51-derived peptides inhibit viral assembly, spread, and accumulation of gE to cell-cell junction. (A) Growth curves for uninfected Vero cells incubated in DMSO or 7nM of indicated peptides. ∼5000 cells were seeded in each well of 96-well plates on day 0. Biomass was determined by crystal violet assay at OD_570_. Culture media were changed every two days. Asterisks above each time point indicate statistical significance of one-way ANOVA analysis. Mean OD_570_ values of time points were also analyzed by one-way repeated measures ANOVA, with the P=0.236 for comparisons between treatments. (B-D) Virus production in Vero cells that were infected with indicated virus at m.o.i=5 and then treated with DMSO or peptides added to the culture medium at 1 h.p.i. Cell lysates were harvested 16 h.p.i. Each bar represents the mean of three independent experiments. (E-G) Plaque size assay of indicated viruses was conducted as in Fig 10, except with the presence of DMSO or 7nM of indicated peptides. (H-J) Plaque size assays of UL51-mutant viruses conducted as in (E-G). (K) Representative images showing accumulation of gE and p150Glued to cell-cell junction in the absence or presence of peptide 90125tat. Vero cells were infected with WT HSV-1(F) BAC virus for 16 hr. (L,M) Quantification of junctional localization of gE (L) and p150Glued (M) performed as described for Fig 10. Scale bars represent 10µm. *=P<0.05, **=P<0.01, ***=P<0.001, ****=P<0.0001. Data represent mean ± SD.

**Table 1.** Sequences of peptides used in Fig 12. Directionalities of peptides are noted by amine groups and carboxylic groups. All peptides contain no modifications.

| Peptide name | Sequence |
| --- | --- |
| 90125 | NH<sub>2</sub>-PGLEAPTIDGAVA<br>AHQDKMRRRLADTCMATILQMYMS-COOH |
| tat90125 | NH<sub>2</sub>-YGRKKRRQRRR PGLEAPTIDGAVAAH<br>QDKMRRRLADTCMATILQMYMS-COOH |
| 90125tat | NH<sub>2</sub>-<br>PGLEAPTIDGAVAAH QDKMRRRLADTCMATILQMYMS Y<br>GRKKRRQRRR-COOH |

### p150Glued a.a 548-911 interacts with pUL51

The sequence in p150Glued that mediates the interaction with pUL51 was also investigated by co-expressing pUL51 with various p150Glued truncations in Vero cells. As shown in Fig 14, B-H, expression of pUL51 with EGFP-p150Glued^1-911^ resulted in filamentous localization of pUL51, while no filamentous pUL51 was detected in cells that expressed EGFP-p150Glued^1-547^. Also, EGFP-p150Glued^548-911^ was enriched on Golgi membranes when co-expressed with pUL51 (Fig 13, G and E), and overexpression of EGFP-p150Glued^548-911^ resulted in a viral growth defect in cells infected with wild-type virus, but not the UL51^d73-244^ mutant (Fig 13, I and J). These results suggest that p150Glued a.a 548-911 plays roles in the interaction with pUL51.

**Fig 13.**
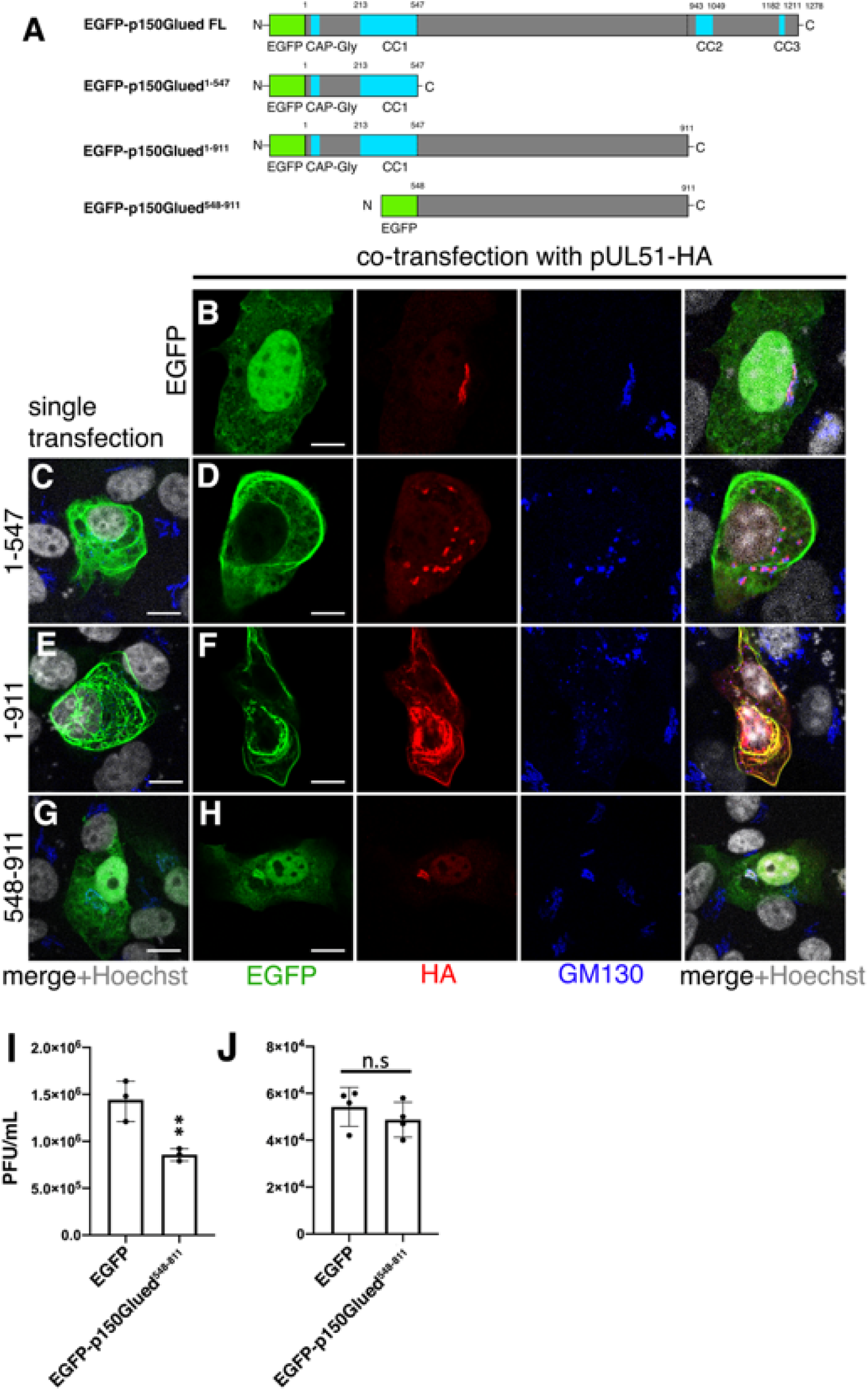
Mapping of p150Glued sequences important for pUL51 recruitment. (A) Schematics depiction of EGFP-p150Glued truncations. Green indicates the EGFP fused to the N-terminus of each construct. Blue indicates known motifs or secondary structures in p150Glued. (B-H) Representative images of Vero cells that were transfected with EGFP-p150Glued mutants only (C, E, G) or co-transfected with pUL51-HA (B, D, F, H). Scale bars represent 10µm. (I, J) Virus production in CAD cells that were transfected with EGFP control or EGFP-p150Glued^548-911^ plasmids 24 hr before infection with WT (I) or UL51^d73-244^ (J) virus for 16 hr. **=P<0.01. Data represent mean ± SD.

**Fig 14.**
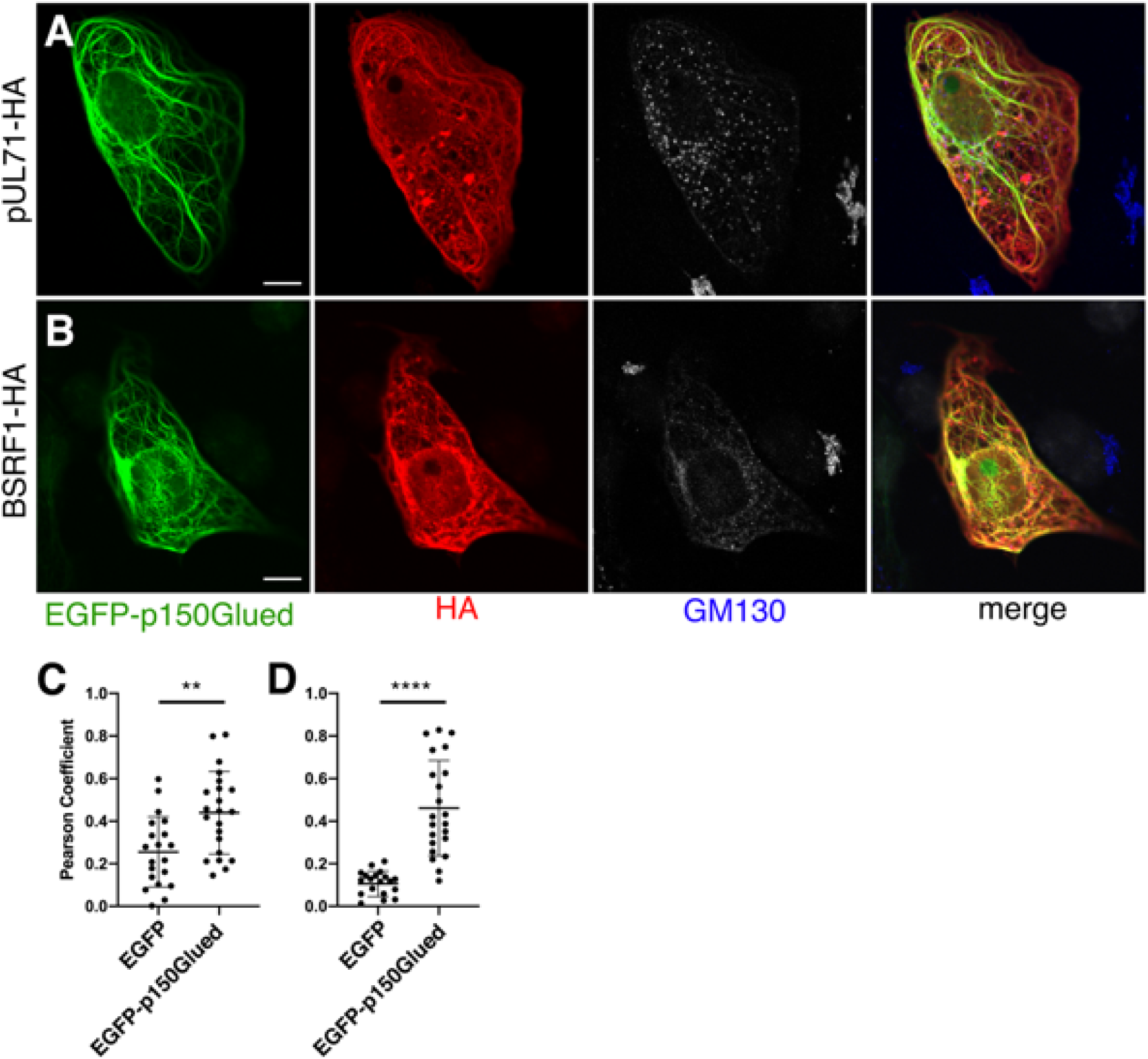
p150Glued recruits pUL51 homologs to microtubule bundles. (A, B) Representative confocal images of Vero cells co-transfected with plasmids that express EGFP-p150Glued and the indicated HA-tagged pUL51 homologs. Scale bars represent 10µm. GM130 split channel is colored in grey. (C, D) Pearson co-efficient for co-localization of EGFP or EGFP-p150Glued with pUL71 (C) or BSRF1 (D) in co-transfected Vero cells. **=P<0.01, ****=P<0.0001. Data represent mean ± SD.

### p150Glued-mediated recruitment of pUL51 homologs

The interaction between p150Glued and pUL51 homolog BSRF1 (EBV) and pUL71 (HCMV) were investigated in co-transfected Vero cells. As shown in Fig 14, both BSRA1 and pUL71 became filamentous in the presence of overexpressed EGFP-p150Glued, suggesting this interaction might be conserved among all families of herpesviruses.

## Discussion

The cytoplasmic assembly of herpesviruses requires that viral structural components, including capsids with associated tegument proteins, vesicles that incorporate viral envelope glycoproteins, and membrane-associated tegument proteins to be brought together in the infected cell cytoplasm. All of these structures are large and might be expected to require cytoplasmic motor function in order to come together efficiently. Indeed, the microtubule-dependency of capsid transportation in HSV is well-established (11, 42, 47). The movement of vesicular cargos, including ER-Golgi transport and retrieval of endosomes from the cell periphery are also dynein-dependent and there is strong evidence that both of these processes are required for efficient HSV assembly (13, 21–23, 48–56).

We have previously shown that HSV organizes its structural components by forming a discrete cVAC in cells of neuronal origin (35). The location of the cVAC, centered on the MTOC, would suggest that minus-end-directed, dynein-mediated transport along microtubules would be important for both cVAC formation and for HSV-1 assembly, and we have previously shown that an intact microtubule network is necessary for cVAC formation. Minus-end-directed microtubule transport requires interaction between the dynein motor and the dynactin complex which serves both as a cargo adapter and as a processivity factor (57). Here, we demonstrate that dynein-dynactin interaction is critical for both cVAC formation and for HSV-1 assembly, since a dominant-negative inhibitor of the interaction disrupts both processes. This inhibition of virus assembly is not secondary to a previously demonstrated function of dynein in facilitating the initiation of viral infection, since we observe the inhibition of viral growth, even when expression of the dominant-negative inhibitor is delayed until after the onset of infection.

Dynein-dependent transport might facilitate movement toward or retention of both membrane vesicles and capsids at the assembly site. While our data clearly show that vesicular transport is disrupted, the effect on capsid movement is less clear. Disruption of dynactin function did not alter the number of capsids that escaped from the nucleus or cause their accumulation at the nuclear rim, but it is possible that proper trafficking of cytoplasmic capsids towards the assembly site is also dynein-dependent and was inhibited by CC1 overexpression.

Dynein-dynactin complexes can have functions other than movement along microtubules, including anchoring microtubules at the MTOC and at the cell periphery (40, 58). Additionally, dynactin can also undergo plus-end-directed, kinesin-dependent transport along microtubules, and is necessary for efficient anterograde transport of some kinesin cargoes (59–61). We observed distinct populations of dynactin that might correspond to these distinct functions. Specifically, we observed two populations of dynactin in infected CAD cells that could be distinguished by their localization in relationship to the cVAC. Dynactin outside the cVAC was localized in small intense puncta without detectable dynein intermediate chain (Fig 2), suggesting that the structures associated with these puncta were not undergoing dynein-dependent, minus end-directed transport. The identity and biological significance of peripheral p150Glued puncta warrant future investigation. One possibility is that these puncta represent vesicles that move away from the cVAC, as many of them also contained gE, pUL11, and capsids. The second population of dynactin was concentrated at the cVAC, and co-localized with dynein intermediate chain (Fig 2). This population of dynactin also appeared to associate with membranes, as its localization pattern (appearing as patches instead of puncta) was the same as that of pUL11, which is distinct from their centrosomal localization in uninfected cells (Fig 2). Since Golgi membranes may serve as a MTOC in HCMV and HSV infected cells (62–64), this population of dynactin may form complexes with dynein for anchoring or nucleation of microtubules at the Golgi (40, 65). Both small puncta and cVAC patches of dynactin remain associated with membranes after nocodazole treatment, unlike centrosome-associated dynactin in uninfected cells (Fig 3), suggesting that the membrane-association of dynactin in infected cells occurs independently of microtubule filaments and may be mediated by one or more viral proteins.

pUL51 is one of the proteins that mediates the interaction between dynactin and membranes that contain virus structural proteins, since it can be recruited by overexpressed p150Glued in infected and co-transfected cells to microtubule bundles, and reciprocally, endogenous cytoplasmic p150Glued can be recruited by overexpressed pUL51 (Fig 4 and 5). The recruitment of pUL51 to overexpressed p150Glued is independent of membrane association of pUL51 or microtubule polymerization, suggesting that recruitment is mediated by formation of a protein complex. The interaction between pUL51 and p150Glued is mediated by a sequence region between a.a 90-166 of pUL51. The N-terminal half (a.a.90-125) of this interaction region evidently provides most of the binding affinity, since it is sufficient for interaction with p150Glued. The C-terminal half (a.a 90-166), by contrast, is insufficient to mediate proper recruitment but seems to enhance the interaction efficiency between pUL51 and p150Glued. The interaction region on pUL51 does not overlap with pUL51 sequences that mediate interaction with its well-established interaction partner pUL7 (39, 66), but instead appears to be in a region that is potentially exposed to the environment (Fig 7). It is possible that pUL51, pUL7, and dynactin could be the same complex in infected cells.

Given that pUL51 mediates interaction with dynactin and the best characterized function of the dynactin complex is to facilitate minus-end directed dynein motor transport, we had expected that disruption of the pUL51-dynactin interaction would inhibit or prevent concentration of pUL51-containing membranes around the centrosome. Indeed, we have previously described a mutation in pUL51 (pUL51^Y19A^) that results in failure of membranes containing viral structural proteins to concentrate around the centrosome (35). However, the properties of the UL51^d90-125^ mutant virus revealed an unexpected level of organization within the HSV cVAC in CAD cells and suggested a different function for pUL51 in organizing the cVAC. In cells infected with the pUL51^d90-125^ mutant virus, membranes containing virus structural proteins still concentrate around the centrosome, but the co-localization between pUL51 and gE (and perhaps other virus structural proteins) is diminished. These results suggest that both concentration of membranes around the centrosome and integration of membranes within the cVAC are mediated by pUL51. Surprisingly, the pUL51-dynactin interaction mediates the latter function rather than the former. Whether pUL51Y19-mediated cVAC formation/maintenance function and a.a 90-125 mediated cVAC formation/maintenance function are in the same pathway warrants further investigation.

pUL51 has previously been shown to have cell-specific roles in promoting both virus spread between cells and virus export to the medium in Vero cells (33) and we show here that both of these functions depend on the dynactin-interaction region of pUL51, since the UL51^d90-125^ mutation diminishes both (Fig 10). This suggests the possibility that pUL51 and dynactin promote trafficking of vesicles containing mature virions away from the cVAC and toward the cell periphery. There are at least three possible ways that pUL51 might mediate this function. First, pUL51 interaction with p150Glued might inhibit its ability to interact productively with dynein and thereby release vesicles that display pUL51 from minus end-directed transport and license them for kinesin-mediated transport toward the cell periphery (Fig 15) (57). This function of pUL51 is regulated by sequences outside a.a 90-125, thus the peptide 90125tat did not disrupt global dynein activity to result in cytotoxicity. Second, pUL51 might alter dynactin function so that it binds to kinesin to mediate plus-end-directed microtubule transport (67). Third, pUL51-dynactin interaction only mediates a short-distance trafficking, which is important to transport mature virion towards a sorting sub-compartment within the cVAC. Since the HCMV cVAC is layered (68), it is possible that HSV cVAC also requires proper arrangement of membranes inside the cVAC for efficient viral egress. It has been recently described that naked capsids can shift from using dynein to using kinesin after arrival at the MTOC during entry via an unknown mechanism (69). It is possible that all microtubule-associated motors are regulated by several tegument or capsid proteins to fine-tune their activities.

**Fig 15.**
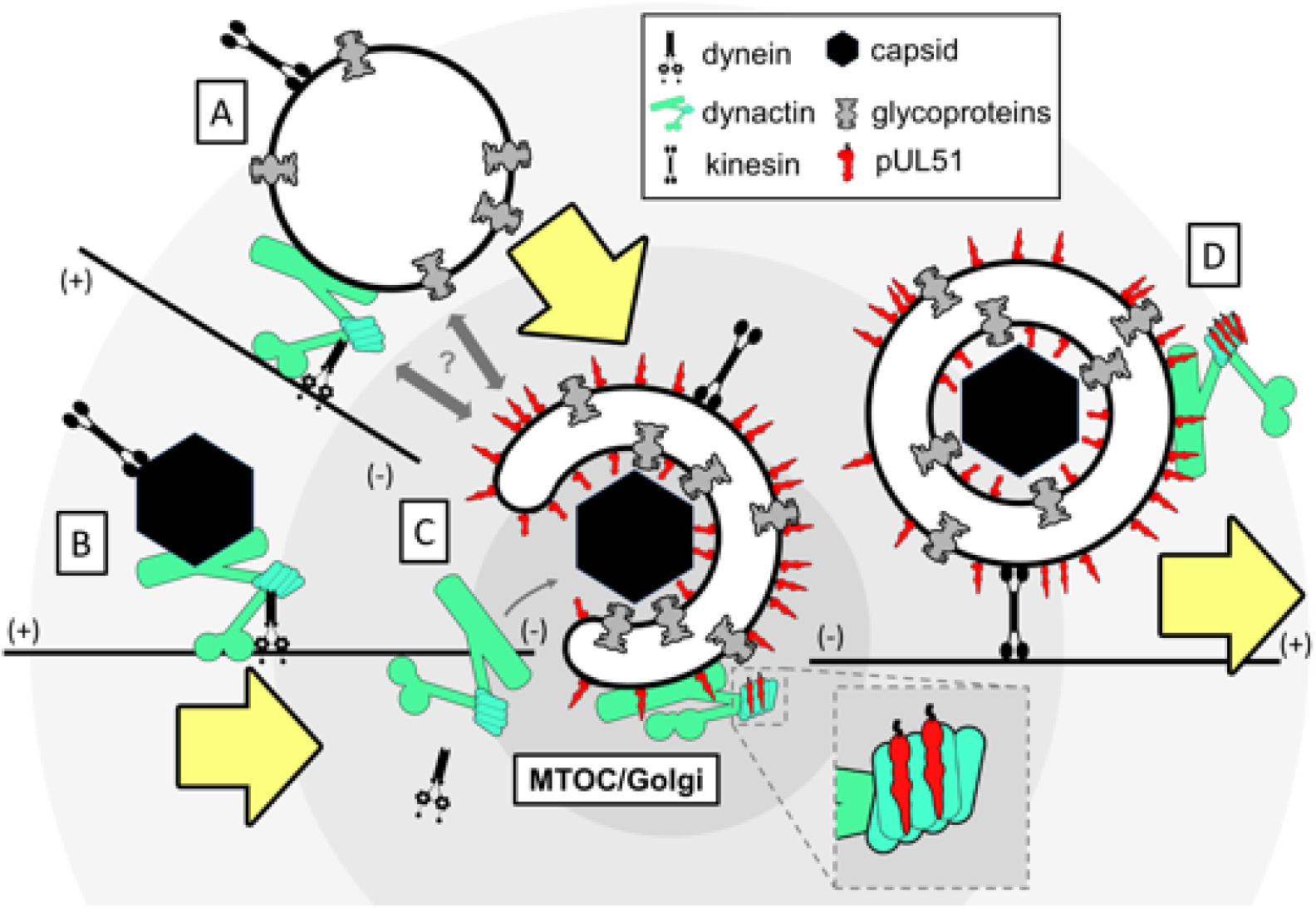
Schematic of a potential mechanism of pUL51-p150Glued interaction. Vesicles containing viral glycoproteins (A) and viral capsids (B) are transported to the cVAC region via dynein-dependent mechanism. (C) after the docking of capsids onto Golgi membranes, pUL51 inhibits dynein function by binding to the elbow region of dynactin, which makes kinesin the preferred motor for vesicle trafficking. (D) kinesin transports mature virions towards cell-cell junction for cell-cell spread.

Cell-to-cell spread of HSV in epithelial cells is thought to be mediated by trafficking of newly produced infectious virions to the junctional surfaces of cells where released virions are sterically protected from neutralizing antibodies and other immune effectors (24). Both pUL51 and gE have well-established roles in promoting spread in epithelial cells (33, 70, 71). Their functions are related, since efficient cell-to-cell spread depends on gE localization to cellular junctions and the expression of pUL51 is necessary for that localization (13, 27, 70, 71). We observed here that p150Glued, but not dynein motors also concentrate at the junctional surfaces of Vero cells in a pUL51-dependent manner. It is possible that dynactin may also facilitate cell-to-cell spread at cell-cell junctions.

Virus assembly pathways are attractive targets for antiviral therapy because they rely on specific interactions between viral factors that can, in principle, be targeted without disrupting host functions. Agents that disrupt this type of interaction, however, select for resistance by mutation in one or both of the interactors. Targeting interactions between virus proteins and host proteins should diminish the potential for development of resistance, since one of the partners is not subject to selection. The 90-125 peptide described here presumably binds to and sterically occludes a binding surface on p150Glued, thereby inhibiting virus spread at very low concentrations. Furthermore, the sequence in p150Glued that is important for pUL51-p150Glued interaction (a.a 548-911) corresponds to a globular region with no known function (Fig 13) (72, 73). This suggests that the 90-125tat peptide might be very useful as a topical agent for inhibition of HSV spread in vivo with high efficacy, low toxicity, and low propensity for development of resistance. Furthermore, the observation that the UL51-p150Glued interaction might be conserved among all herpesviruses suggests the exciting possibility that agents that disrupt this interaction might show broad efficacy in treating herpesvirus infections.

## Materials and Methods

### Cell lines

CAD cells (gifts from David Johnson at the Oregon Health & Science University) were maintained in Dulbecco modified Eagle medium (DMEM) F-12 (Thermo Fisher Scientific) supplemented with 10% fetal bovine serum (FBS) and penicillin-streptomycin (P/S). SH-SY5Y cells (gift from Stevens Lewis at the University of Iowa) were maintained in Opti-MEM supplemented with sodium pyruvate, non-essential amino acid, P/S, and 10% FBS. Vero and HEp2 cells (purchased from ATCC) were maintained in DMEM supplemented with 5% bovine caff serum and P/S. HSV pUL51-complementing cells were constructed before (33), and were maintained in DMEM supplemented with 5% FBS, P/S, and 0.3 mg/ml G418. 293T cells (gifts from Mark Stinski at the University of Iowa) were maintained in DMEM supplemented with 10% FBS and P/S. All cells were cultured in a humidified incubator at 37°C with 5% CO2.

### Plasmids

pEGFP-p150Glued was acquired from Addgene (#36154). pcDNA-BSRF1-HA was custom made using GeneArt service from Thermo-Fisher. pcDNA3-pUL51-FLAG full length and truncation plasmids were generated before (27). All other plasmids generated in this study were assembled by Gibson Assembly Master Mix (New England Biolabs Inc.) according to manufacturer instruction. For pcDNA3-pUL51-HA plasmid, the pUL51 fragment was amplified from HSV-1(F) BAC using primers UL51HAfwd and UL51HArev, and backbone fragments were amplified from pcDNA3 with primer pairs vector1fwd + vector1rev and vector2fwd + vector2rev. For pcDNA3-pUL51^C9V^-HA plasmids, two backbone fragments were amplified from pcDNA3-pUL51HA using primer pair CDNA7 + UL51seq7 and CDNA8 + UL51seq6, and pUL51^C9V^ insertion was amplified from an HSV-1(F) pUL51^C9V^ virus using primers UL51seq5.5 + UL51seq8. For pcDNA3-pUL51^Y19A^-HA, pcDNA3-pUL51^I97A^-HA, pcDNA3-pUL51^R109A, R110A^-HA, pcDNA3-pUL51^L111A^-HA, pcDNA3-pUL51^G91A, L92A, E93A^-HA, pcDNA3-pUL51^d90-125^-HA plasmids, mutations were introduced by amplifying pcDNA3-pUL51-HA with the following primer pairs respectively: Y19A fwd + CDNA8, CDNA7 + Y19A rev; 2R2A fwd + CDNA8, CDNA7 + 2R2A rev; I97A fwd + CDNA8, CDNA7 + I97A rev; L111A fwd + CDNA8, CDNA7 + L111A rev; AAA fwd + CDNA8, CDNA7 + AAA rev; CDNA37 + CDNA7, CDNA38 + CDNA8. Fragments for pEGFP-CC1 plasmid were amplified from pEGFP-p150Glued using primers DNCT1, DNCT2, DNCT3, DNCT4, DNCT7, and DNCT8. For 34p-EGFP and 34p-CC1 plasmids, the 388-bp sequence before the natural start codon of HSV UL34 gene was amplified from HSV-1(F) BAC using primers pHSV1 + pHSV2. Other fragments for this plasmid were amplified from pEGFP C1 (Promega) or pEGFP-CC1 using primer GFP3, CDNA7, CNDA8, and CDNA9. For the 34p-EGFP-IRES-VP5 plasmid, the IRES fragment was amplified from pTUNER-IRES (Promega) with primers IRES10 + IRES11, the VP5 gene was amplified from HSV-1(F) BAC with primer pairs IRES12 + VP5midRev and VP5midFwd + IRES15, the EGFP fragment was amplified from 34p-EGFP with primers pHSV1 + IRES14, the backbone fragment was amplified from 34p-EGFP with primers CDNA28 + CDNA9. For the 34p-CC1-IRES-VP5 plasmid, the IRES fragment was amplified from pTUNER-IRES (Promega) with primers IRES10 + IRES11, the VP5 gene was amplified from HSV-1(F) BAC with primer pairs IRES12 + VP5midRev and VP5midFwd + IRES13, the CC1 fragment was amplified from 34p-CC1 with primers pHSV1 + IRES9, the backbone fragment was amplified from 34p-CC1 with primers DNCT3 + CDNA9. For the pcDNA3-pUL51^90-125^-mCherry plasmid, pUL51^90-125^-mCherry fragment was amplified from pLVX-IRES-mCherry (gift from Balaji Manicassamy at the University of Iowa) using primers RFP3 + CDNA17 (which adds pUL51^90-125^ sequence to mCherry), which was then extended with primers RFP4 + CDNA17, backbone fragments were amplified from pcDNA3-pUL51-HA using primers CDNA7 + RFP5 and IRES5 + CDNA8, with the former fragment then extended with primers CDNA7 + RFP6. For the pcDNA3-mCherry plasmid, the mCherry fragment was amplified from pLVX-IRES-mCherry using primers CDNA33 + CDNA34, the backbone fragments were amplified from pcDNA3 with primers CDNA1 + CDNA2 and CDNA3 + CDNA4. For pEGFP-p150Glued^1-547^ and pEGFP-p150Glued^1-911^ plasmids, backbone fragments were amplified from pEGFP-p150Glued using primer pairs DNCT1 + DNCT2 and DNCT3 + DNCT4, insertion were amplified from pEGFP-p150Glued using primer pairs DNCT9 + DNCT8 or DNCT9 + DNCT13. For pEGFP-p150Glued^548-911^ plasmid, fragments were amplified from pEGFP-p150Glued^1-911^ plasmid using primers DNCT1 + DNCT14 (extended with DNCT1+ DNCT15) and DNCT16 + DNCT4. For FLAG-EGFP-p150Glued and FLAG-EGFP-p150Glued^1-911^ plasmids, four fragments were amplified from respective untagged plasmid using primer pairs GFP5 + DNCT18, DNCT17 + SV40 polyA rev, SV40 polyA fwd + DNCT4, and DNCT1 + GFP6. For FLAG-EGFP, FLAG-EGFP-p150Glued^1-547^, and FLAG-EGFP-p150Glued^548-911^ plasmids, three fragments were amplified from respective untagged plasmid using primer pairs GFP5 + SV40 polyA rev, SV40 polyA fwd + DNCT4, and DNCT1 + GFP6. For pcDNA3-pUL71-HA, the pUL71 fragment was amplified from HCMV-infected cell lysate (kindly provided by Jeff Meier at the University of Iowa) using primer pair UL71HAfwd + UL71HArev, then extended with primers UL71HAfwd + CterHAfwd. All plasmids were sequenced to confirm promoter and insertion sequences. All primer sequences are listed in Supplemental Table 1.

### HSV-1(F) BACs

HSV-1(F) BAC was a gift from Yasushi Kawaguchi at the University of Tokyo. The HSV-1(F) VP5-null virus was a gift from Prashant Desai at Johns Hopkins University. HSV-1(F) UL35-RFP BAC virus was a gift from Cornel Fraefel at the University of Zürich. HSV-1(F) pUL51-HA and HSV-1(F) pUL51^Y19A^-HA virus was described before (White 2020). pUL51^C9V^ and pUL51^d90-125^ mutations were introduced in HSV-1(F) or HSV-1(F) pUL51-HA backbone respectively using en passant protocol (74). Details and primer sequences are described in Supplemental Table 2. Mutations in viral stock were confirmed by PCR amplifying UL51 region use primer UL51seq1 and UL51seq3 followed by Sanger sequencing using primer UL51seq1.

### Peptides

Chemically synthesized recombinant peptide 90125, tat90125, 90125tat were purchased from Alan Scientific Inc. Lyophilized peptide power was reconstituted in DMSO (Sigma) to 0.7 mM as stock solutions.

### Transfection of cells

For immunofluorescent microscopy, 1×10^5^ cells of CAD, SH-SY5Y, Vero, or HEp2 cells were seeded into 24-well tissue culture plates with a 13-mm-diameter coverslips in each well 17-24 hr before transfection. CAD and Vero cells were then transfected with X-tremeGENE™ HP DNA Transfection Reagent (Millipore-Sigma). SH-SY5Y and HEp2 cells were transfected with Lipofectamine® Reagent (Invitrogen). All transfections followed product protocols. For co-IP, 4×10^6^ 293T cells were seeded in 100mm plates 24 hr before transfection. 3 μg of DNA and 72μg of poly-ethylenimine (diluted to 1 μg/μl stock solution in ddH_2_O) were mixed in 500 μl of RPMI (Gibco), and incubated 15 min before adding to each plate containing 20ml of 293T cell medium.

### Infection and viral growth assay

For all infections, viruses were diluted in infection medium (DMEM containing 1% heat-inactivated serum and P/S) to achieve m.o.i of 5. Viruses were incubated with cells for 1hr before the removal of the inoculum. Residual viruses from the inoculum were further neutralized with a sodium citrate buffer (50 mM sodium citrate, 4 mM KCl, adjusted to pH=3 with HCl), after which cells were washed with the infection medium for two times before adding 1ml of infection medium. For all transfection-superinfections, cells were transfected 24 hr before infection. For DMSO or peptide treatment, DMSO or peptide DMSO solutions were added in the replacement infection medium.

For viral growth assays, the concentration of infectious virions was titrated on monolayers of pUL51-complementing cells by plaque assay. For viral release, the supernatant of each well of cells was harvested first, then 1ml fresh infection medium was added to resuspend all infected cells. Harvested samples were first frozen, thawed, and then sonicated before titration.

### Immunofluorescence assays

Transfected or infected cells on 13-mm-diameter coverslips were fixed with 3.7% formaldehyde for 20 min before washing with PBS two times. The immunofluorescent (IF) buffer used to dilute antibodies or antiserum is a PBS buffer that contains 1% triton X100, 0.5% sodium deoxycholate, 1% egg albumin, and 0.01% sodium azide. Fixed coverslips were first blocked with IF buffer containing 10% human serum for 30min before 1 hr of primary antibody staining (with 10% human serum) and 1 hr of secondary fluorescent antibody staining. The primary antibodies were diluted as follow: rabbit anti-HA epitope tag antibodies (Abcam) 1:500, rabbit anti-UL51 antiserum (gift from Joel Baines at the Cornell University) 1:500, rabbit anti-UL11 antiserum (gift form John Wills at the Penn State University) 1:500, mouse (DL6) anti-gD monoclonal antibody (ATCC) 1:1,500, mouse polyclonal anti-gE antibody (Virusys) 1:500, goat anti-p150Glued antibody (Abcam) 1:500, mouse anti-DIC antibody (Santa Cruz Biotechnology) 1:500, rabbit anti-Rab11 antibody (Abcam) 1:250, mouse anti-GM130 antibody (BD Bioscience) 1:500, mouse anti-ß-catenin antibody (BD Bioscience) 1:500, and 1:500 mouse anti-ß-tubulin antibody (Sigma-Aldrich). All secondary antibodies were diluted 1:1000. Nuclei were stained with 1:500 DNA dye Hoechst 33342 (Invitrogen). Stained coverslips were mounted with ProLong^TM^ Diamond Antifade Mountant (Invitrogen) and let dry for 12 hr. Confocal images were taken on a Leica DFC7000T confocal microscope equipped with Leica software.

For Pearson co-efficient measurement for protein co-localization, confocal images of 1024×1024 pixels were captured for random fields. Pearson co-efficient between signals of two different proteins was measured in ImageJ FIJI for each cell. The boundary of each cell was defined by either gE signals or ubiquitous cytoplasmic fluorescent proteins.

For cytoplasmic capsid quantification, the number of cytoplasmic capsids in infected cells were quantified by using a modified protocol that has been previously described (5). Hoechst 33342 staining and gE staining were used to define the nuclear and cytoplasmic boundaries of the cell, respectively. ImageJ was used to count the number of RFP maxima in the cytoplasm.

### Western blotting

For cell lysate samples, cells from each well of a 6-well plate were pelleted and lysed in P.I. RIPA buffer (50 mM Tris pH=7.5, 150mM NaCl, 5 mM NaF, 5mM sodium vanadate, 1mM EDTA, 1% triton X-100). Lysates were then sonicated for 10 s before adding loading dye (5x stocks contain 4% SDS, 0.25M Tris buffer pH=6.8, 0.6mg/ml bromphenol blue, and 30% glycerol) and 2-mercaptoethanol (to a final concentration of 1%). Proteins were denatured by incubating at 85 °C for 8 min. Denatured samples were then separated on SDS-PAGE by size and transferred to nitrocellulose membranes before blocking in 5% nonfat milk plus 0.2% Tween 20 for at least 1 hr. Then, membranes were probed with either 1:500 rabbit anti-UL51 antiserum, 1:500 mouse anti-VP5 antibody (Biodesign), 1:5,000 rabbit anti-EGFP antiserum (gift from Craig Ellemeier at the University of Iowa), 1:500 rabbit anti-mCherry antiserum (gift from David Weiss at the University of Iowa), or 1:250 chicken anti-UL34 antiserum (75) and visualized by alkaline phosphatase-conjugated secondary antibody.

### Co-immunoprecipitation

100mm plates of transfected 293T cells were lysed in 1ml of P.I. RIPA buffer with SigmaFAST™ protease inhibitor cocktails (Sigma) for 5 min. Nuclei and cellular debris were pelleted and discarded. 8 μl of anti-FLAG magnetic beads (Sigma-Aldrich) was then added to each sample, followed by gentle rotation overnight at 4°C. Beads were then separated magnetically and washed with P.I. RIPA buffer for three times. Proteins on beads were eluted with P.I. RIPA buffer containing FLAG peptide (ApexBio) by incubating on ice for 5 min. Loading dye and 2-mercaptoethanol were then added to samples before denaturing by incubating at 85 °C for 8 min. Samples were then analyzed by Western blotting.

### Plaque size assay

1.8×10^6^ Vero cells were seeded into each well of a 6-well plate to make a cell monolayer the day before infection. The infection was initiated by replacing growth medium with 1ml of infection medium containing 100-1000 plaque-forming units (PFU) of WT HSV or mutants. The inoculum was allowed to incubate for 1.5hr before being replaced with 2.5ml of infection medium containing 1:250 dilution of GammaSTAN S/D (Talecris Biotherapeutics), which contains neutralizing antibody against HSV. Monolayers of cells were then fixed at 48 h.p.i with 3.7% formaldehyde for 20 min before washing with PBS two times. Fixed monolayers of cells were then stained for 1 hr using 1:2,500 dilution of mouse monoclonal anti-HSV 45-kDa protein (scaffolding protein) antibody (Serotec) as a primary antibody, and stained for another 1 hr using 1:1,000 dilution of Alexa Fluor 568 goat anti-mouse IgG (Invitrogen) as a secondary antibody. Images of plaques were taken with an inverted fluorescence microscope. Plaque images were analyzed in ImageJ and outlined by using the freehand tool. The number of pixels contained within the outline was used as the plaque area.

### Cell growth assay

∼5,000 Vero cells were seeded into each well of a 96-wells plate 16 hr before replacing culture medium with medium containing peptides or DMSO. The growth medium was replaced for each well with identical fresh medium every 48 hr. Every 24 hr, cells were washed with PBS, and fixed/stained for 15min with 100 μl staining reagent containing 20% methanol and 0.5% crystal violet. After staining, each well was washed five times with distilled H_2_O. After an air-dry of at least 1 hr, the crystal violet dye in fixed cells was eluted with 100 μl 10% acetic acid. The dye concentration in elution liquid were measured by a Tecan infinite M200Pro plate reader at 570nm. Blank wells contained growth medium but not cells were used as controls.

### Statistical analysis

All statistical analyses were conducted by using student’s T-test, one-way ANOVA, or one-way repeated measures ANOVA in GraphPad Prism 8. The number of cytoplasmic capsid local maxima, pixel number of plaques in plaque size assays, and PFU number in infectious particle titration were log-transformed before statistical analyses.

## Acknowledgments

The authors would like to thank Drs. Joel Baines, John Wills, and Harvey Friedman for providing antisera and Masoudeh Masoud Bahnamiri and Alison Haugo-Crooks for critical reading of the manuscript. This work was funded by NIH grant R21AI153683. S. W. was supported by a University of Iowa Graduate College Fellowship. M.T. was supported by NSF REU DBI-1852070.

## Author Contributions

Shaowen White: Conceptualization, Data Curation, Formal Analysis, Funding Acquisition, Investigation, Methodology, Writing - Original Draft

My Tran: Investigation, Formal Analysis

Richard Roller: Conceptualization, Funding Acquisition, Investigation, Project Administration, Supervision, Writing – review and editing

---

**Fig S1.**
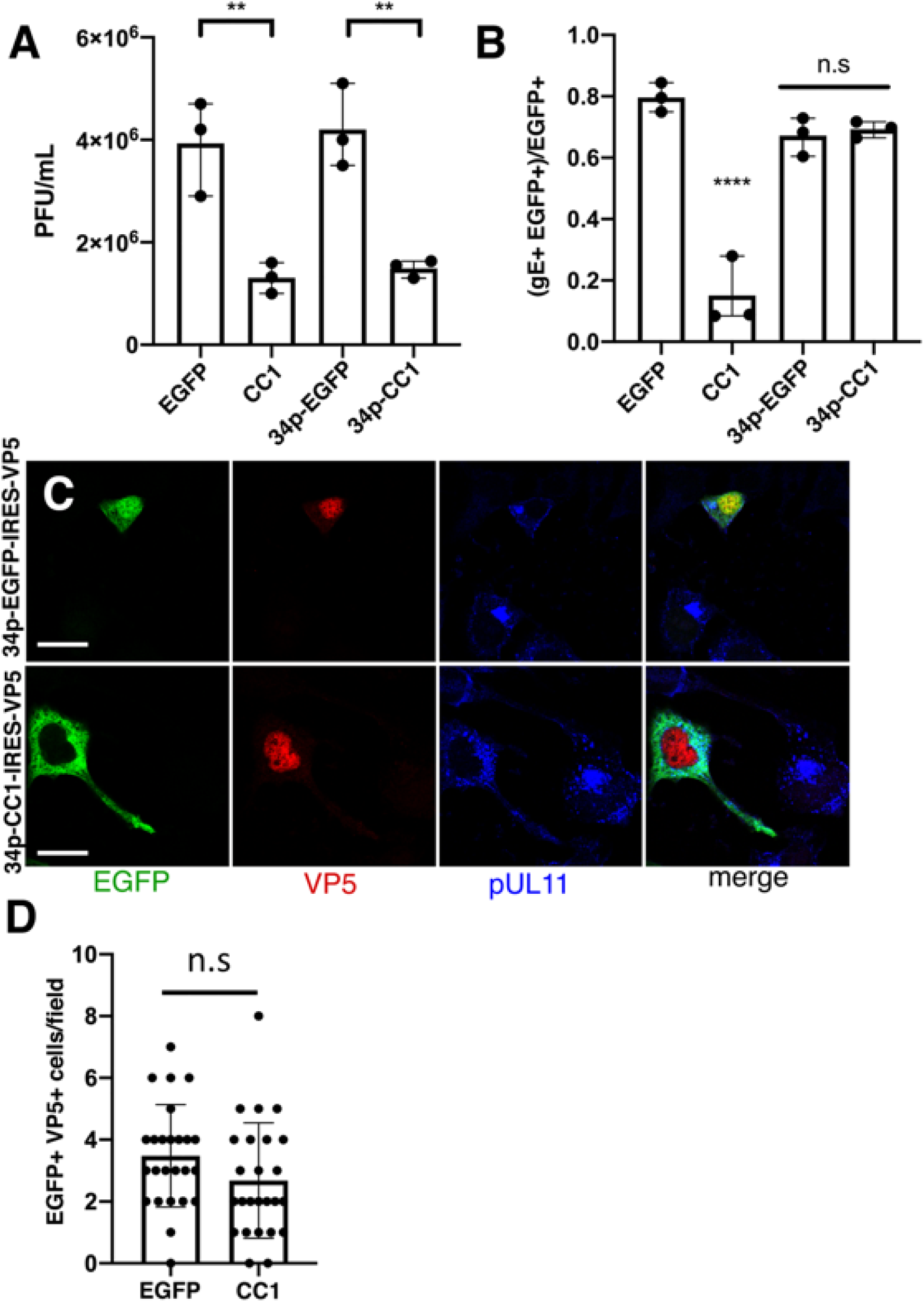
34p-CC1 transfection did not prevent the expression of viral late gene gE. (A) Viral production in transfected-superinfected CAD cells at 16 hr after infection at m.o.i.=5. (B) Percentage of infected CAD cells that expresses GFP or CC1, quantified by confocal imaging. At least 50 GFP-positive cells were imaged and scored for presence of gE from three independent experiments. (C, D) CAD cells were transfected with indicated plasmid before infected with VP5-null virus for 12 hr. (D) The number of EGFP-VP5 double-positive cells were counted for at least 25 random fields. **=P<0.01, ***=P<0.001, ****=P<0.0001. Data represent mean ± RNG.

**Fig S2.**
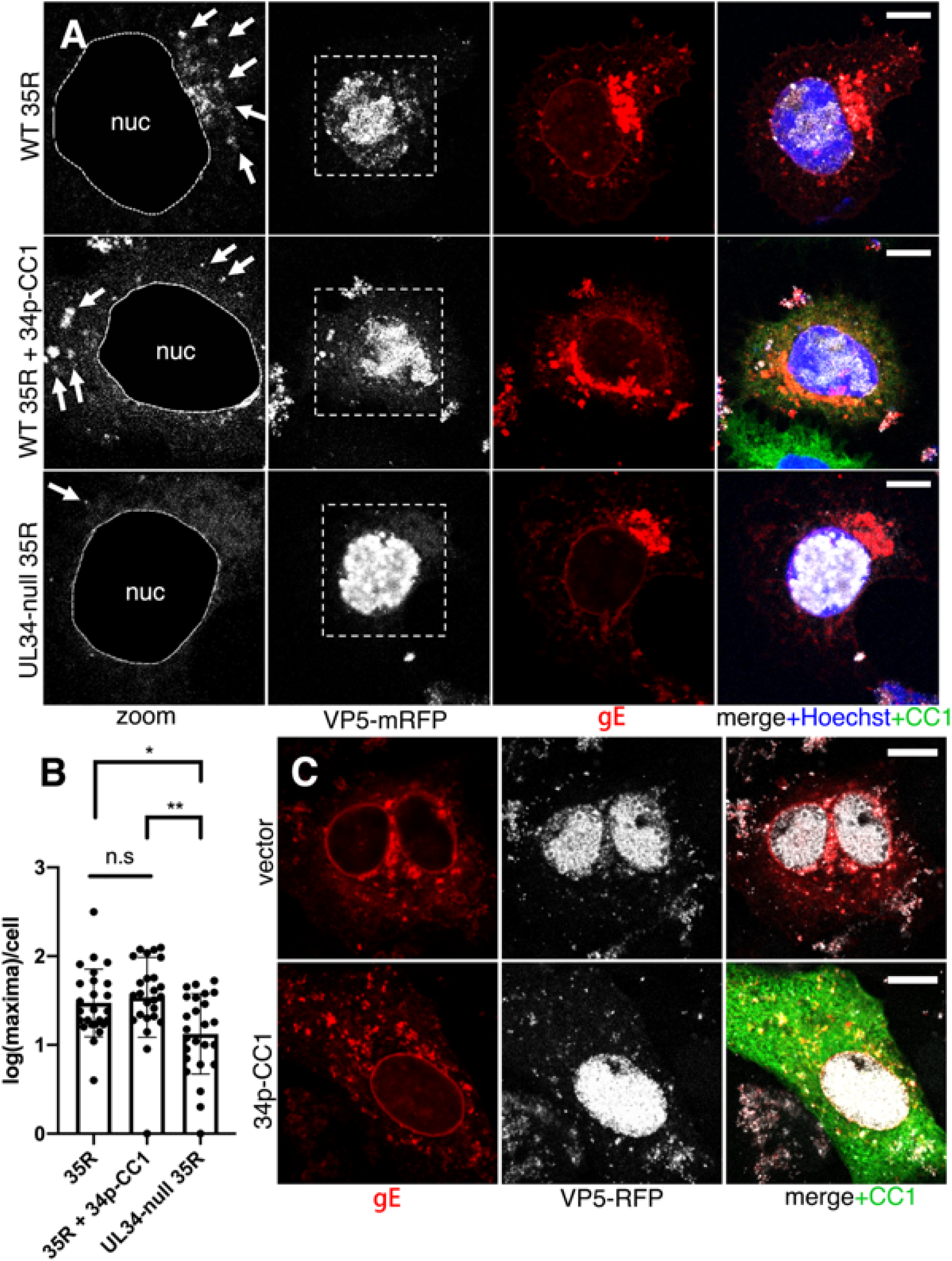
CC1 overexpression does not prevent capsid nuclear egress. (A) Confocal images of CAD cells infected with HSV-1 35R virus, which expresses a mRFP fusion to the capsid protein VP26. Nuclei are stained with Hoechst33342 in blue; CC1-EGFP is shown in green, gE detected with anti-gE antibody is shown in red and mRFP-VP26 is shown in white. In the middle row (CC1 expressing cells), CAD cells were transfected-superinfected as described in Fig 1. In zoomed images in the left-most column, nuclei are masked (guided by Hoechst staining). White arrows indicate individual or aggregated cytoplasmic capsids. (B) Quantification of cytoplasmic capsids per cell by counting local mRFP maxima in the cytoplasm of 25 cells per condition. (C) Same as (A) but in Vero cells. All scale bars represent 10µm. *=P<0.05, **=P<0.01. Data represent mean ± SD.

**Fig S3.**
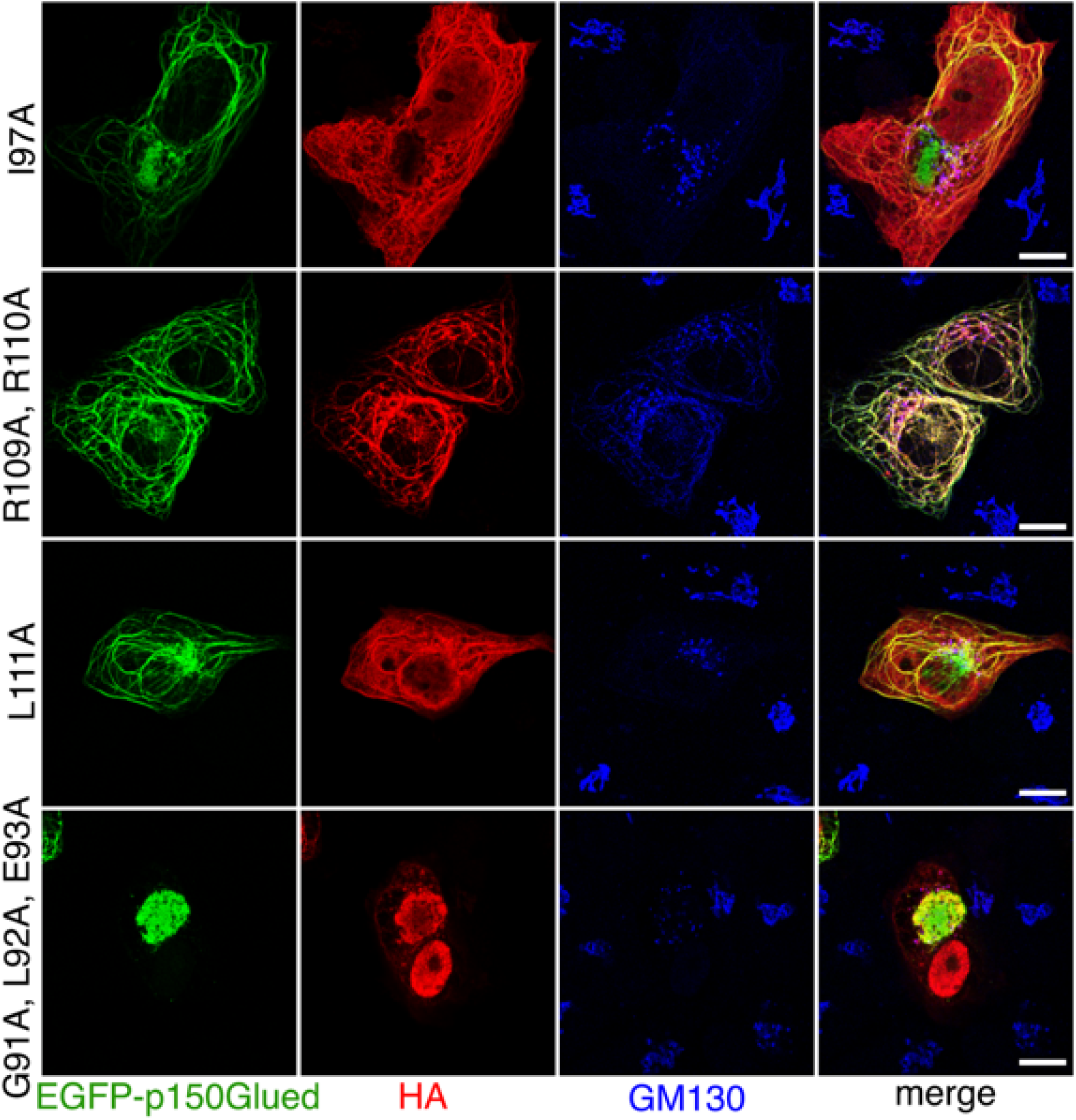
pUL51 mutants that still interacts with p150Glued. Vero cells were co-transfected with pUL51-HA mutants and EGFP-p150Glued. Scale bars represent 10µm.

**Table S1.**
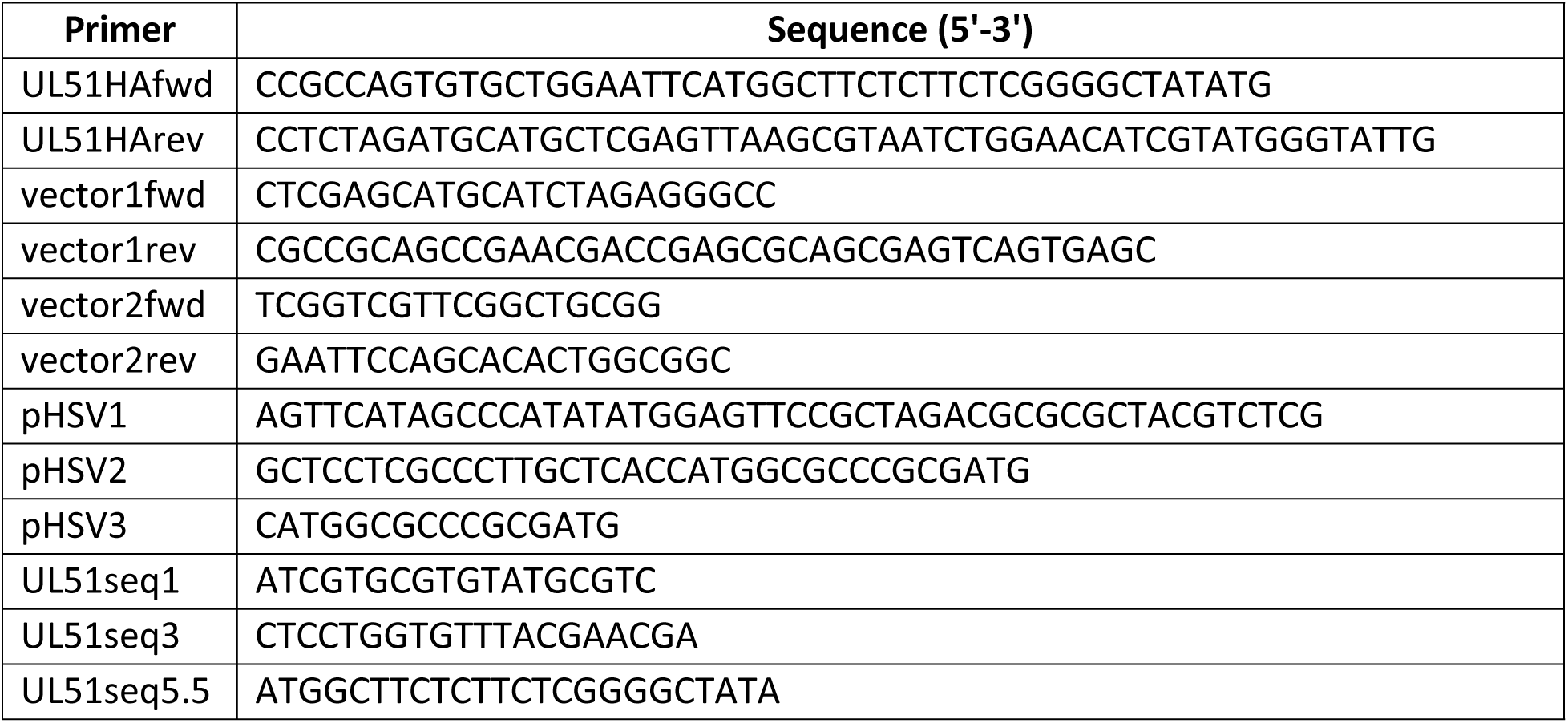
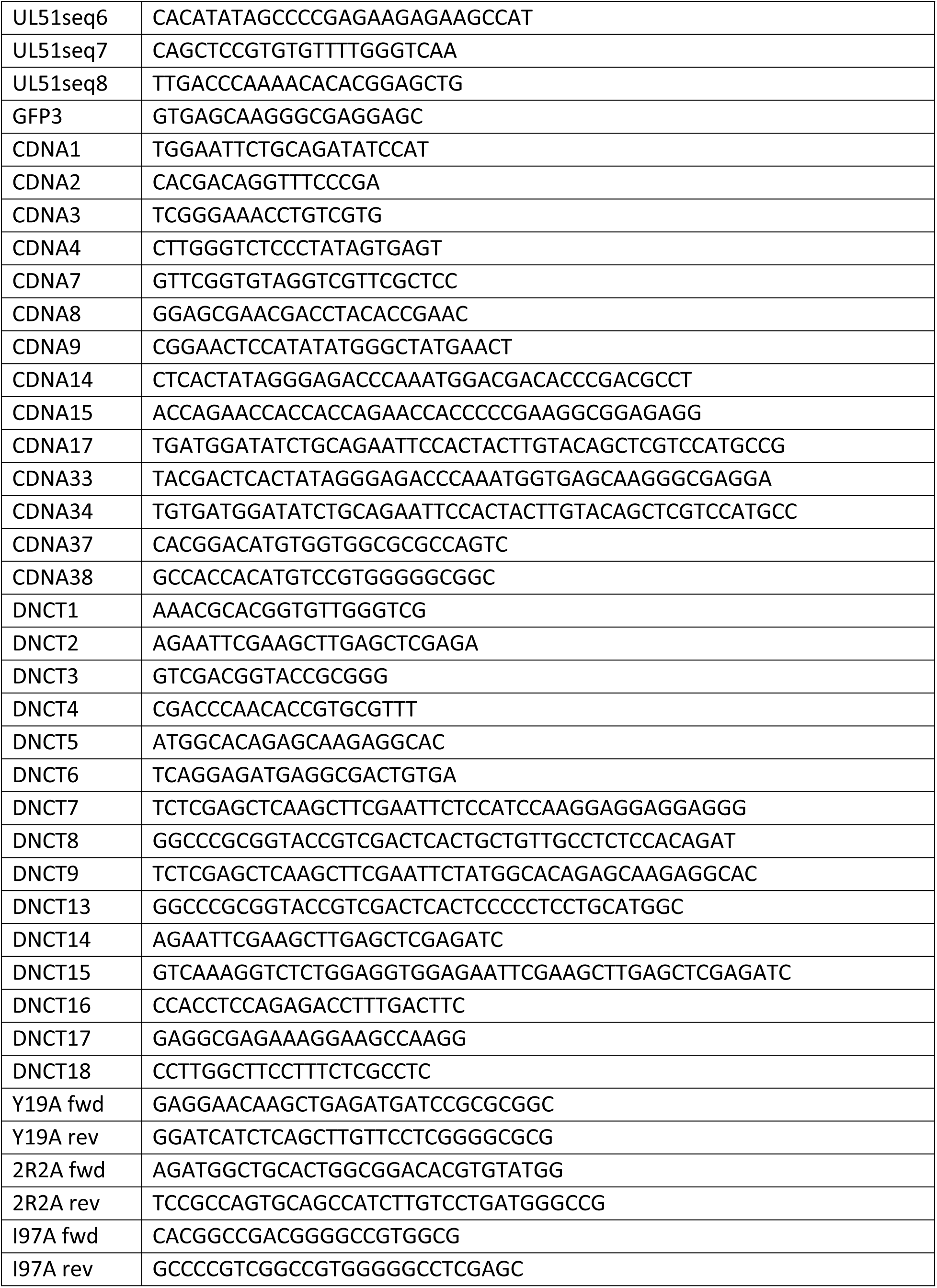
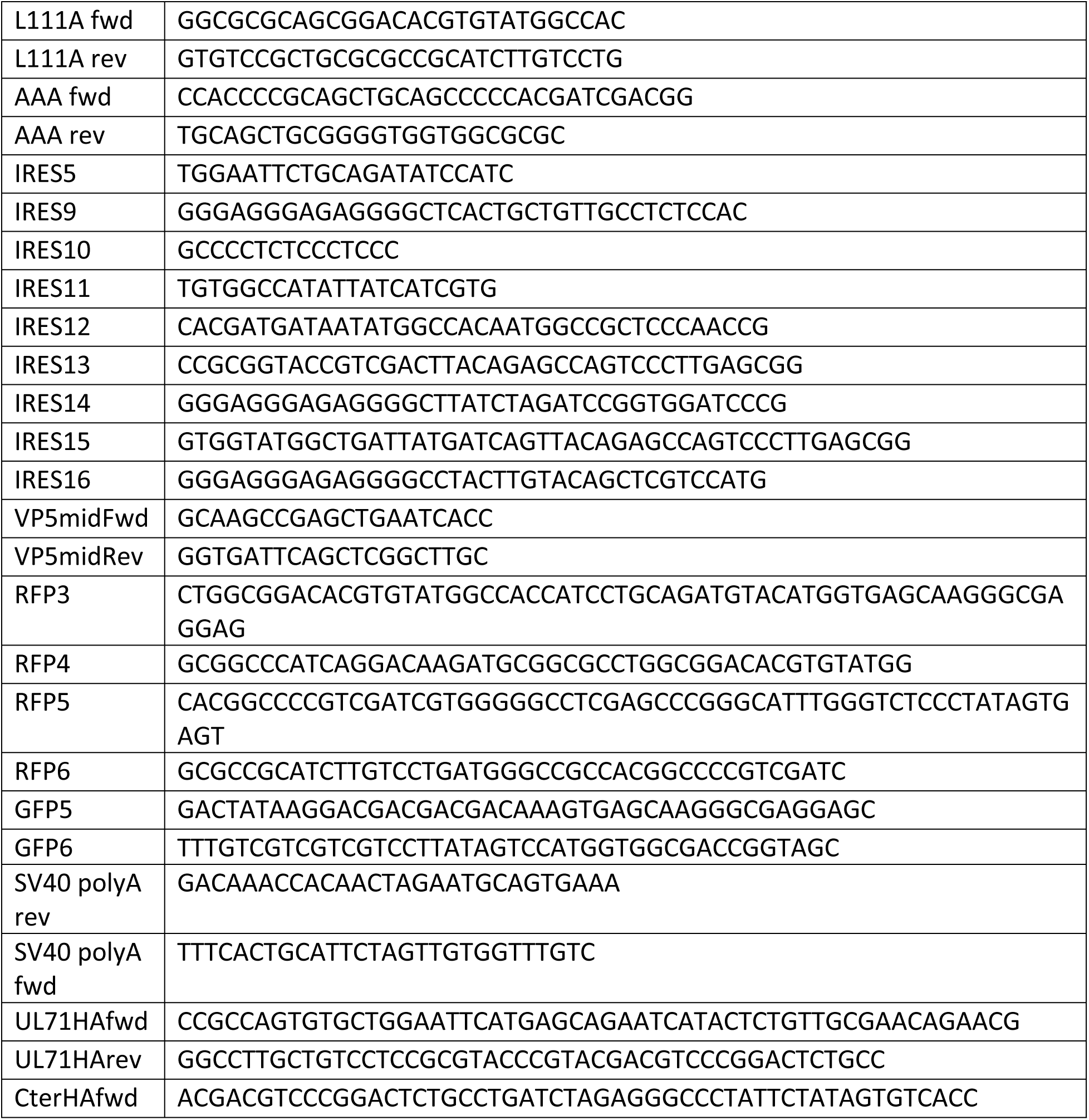
Sequences of primers used in DNA vector cloning. All primers were purchased from Integrated DNA Technologies (Coralville, Iowa).

**Table S2.**
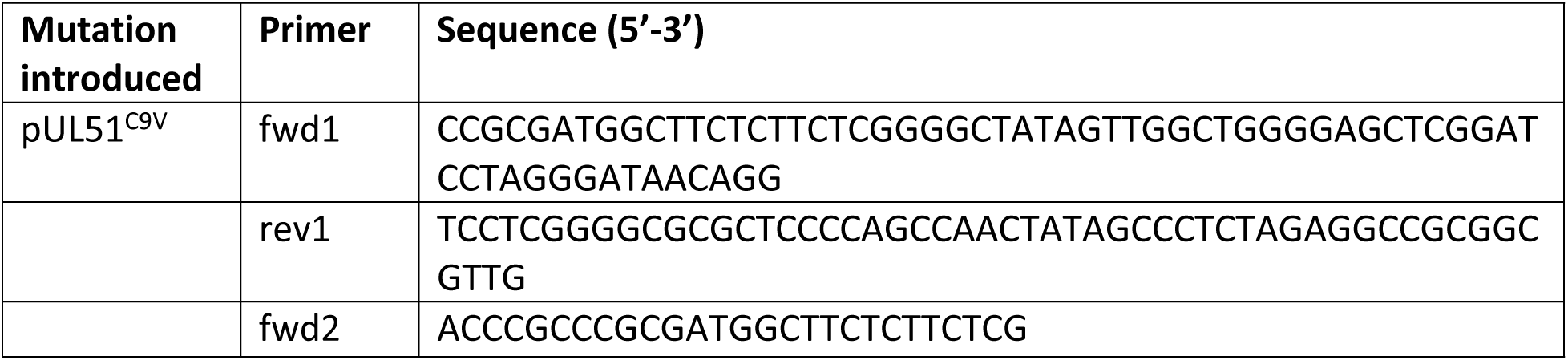
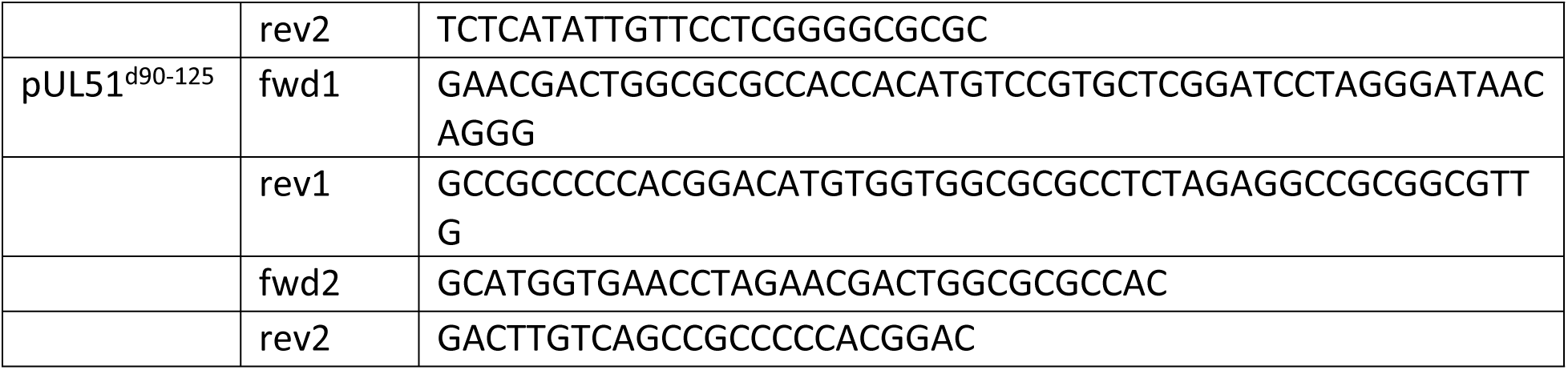
Sequences of primers used to generate mutagenic DNA fragment for HSV-1(F) BAC engineering. For all mutagenic DNA fragments, a gentamicin resistance (GmR) cassette containing a SceI homing nuclease site and pUL51 recombination sequences was constructed as the following: two fragments containing flanking sequences with each other were amplified from pEGFP-SceI-GmR using primer pairs fwd1 + midGentRev (5’-GCGGTTGTTGGCGCTCTCG-3’) and midGentFwd (5’-CACTACGCGGCTGCTCAAACC-3’) + rev1. Two fragments were then recombined into one piece by PCR amplification with outer primers fwd2 and rev2. The mutagenic DNA fragment was then recombined into HSV-1(F) BAC in lambda Red recombinase-expressing GS1783 *E.coli*. Scarless excision of the Gm cassette was induced following *En Passant* mutagenesis protocol.

